# Single-molecule analysis of transcription activation: dynamics of SAGA co-activator recruitment

**DOI:** 10.1101/2023.08.07.552353

**Authors:** Jongcheol Jeon, Larry J. Friedman, Hogyu David Seo, Augustus Adeleke, Bria Graham, Emily Patteson, Jeff Gelles, Stephen Buratowski

## Abstract

Transcription activators are said to stimulate gene expression by “recruiting” coactivators to promoters, yet this term fits several different kinetic models. To directly analyze dynamics of activator-coactivator interactions, single-molecule microscopy was used to image promoter DNA, a transcription activator, and the Spt-Ada-Gcn5 Acetyltransferase (SAGA) complex within nuclear extract. SAGA readily, but transiently, binds nucleosome-free DNA without activator, while chromatin template association occurs nearly exclusively when activator is present. On both templates, activator increases SAGA association rates by up to an order of magnitude, and dramatically extends its dwell times. These effects reflect direct interactions with the transactivation domain, as VP16 or Rap1 activation domains produce different SAGA dynamics. Despite multiple bromodomains, acetyl-CoA or histone H3/H4 tail acetylation only modestly improves SAGA binding. Unexpectedly, histone acetylation more strongly affects activator residence. Our studies thus reveal two modes of SAGA interaction with the genome: a short-lived activator-independent interaction with nucleosome-free DNA, and a state tethered to promoter-bound transcription activators that can last up to several minutes.

## Introduction

Eukaryotic gene expression is typically regulated at the level of mRNA production, most commonly by transcription activators bound to promoter elements known as enhancers or Upstream Activating Sequences (UASs). Eukaryotic transcription activators have distinct domains for DNA binding and activating functions, both of which are needed to bring various coactivator enzymes to target promoters (Hahn and Young, 2011). One major class of coactivators includes the histone acetyltransferases (HATs) and ATP-dependent chromatin remodelers (Cairns, 2009; Li et al., 2007), which remove or shift nucleosomes that occlude the core basal promoter. A second class of coactivators interact directly with basal initiation factors to promote assembly of pre-initiation complexes (PICs) at the core promoter. This group includes the TBP-associated factors (TAFs), which chaperone TATA-Binding Protein (TBP) to promoters as part of the TFIID complex (Patel et al., 2020), and Mediator (Allen and Taatjes, 2015; Malik and Roeder, 2016), which acts as a bridge between transcription activators and RNA polymerase II (RNApII).

Although activators are proposed to “recruit” coactivators, this term is generally not defined kinetically (Karr et al., 2022). Although coactivators can be crosslinked at activator binding sites, chromatin immunoprecipitation levels reveal only steady-state occupancy, not dynamics. Similarly, in vitro binding studies of activator-coactivator interactions are typically done under equilibrium conditions. As first conceptualized to explain transcription activation in prokaryotes (Ptashne, 2005; Ptashne and Gann, 1997), “recruitment” is often used to describe classic cooperative binding. In this mechanism, the regulatory transcription factor and its target each independently bind distinct DNA sequences, with protein-protein contact between them stabilizing the bound state. This interaction primarily manifests as a decrease in the dissociation rates of both proteins, hence the term “cooperativity”. This paradigm is illustrated by prokaryotic transcription activators that contact both DNA and the sigma 70 - RNAP holoenzyme. Stabilizing the short-lived “closed” polymerase complex on the promoter increases the time available for slower promoter melting and conversion to an “open” complex competent for RNA synthesis.

A second “recruitment” model proposes activators tethering coactivators at the UAS/enhancer through protein-protein interactions, increasing both coactivator association rate and the local concentration available for transfer to a nearby promoter or nucleosome. In one version of this model, the coactivator remains tethered to activator throughout its cycle of activity on the template. In the alternative “hit and run” or “kiss and run” variation of this model, once the coactivator transfers to its nearby site of action, activator contact is broken and the activator is no longer required to retain coactivator on the template. Yet another recruitment model invokes activator contact triggering a conformation change in the coactivator to facilitate subsequent functions or interactions, as proposed for Mediator (Zhang et al., 2021), SAGA (Mittal et al., 2018), and p300 (Karr et al., 2022). To date, there have been few careful quantitative kinetic analyses to help distinguish these models.

Here we analyze interactions of DNA-bound activators with the SAGA (Spt-Ada-Gcn5 acetyltransferase) complex. Early yeast genetic studies implicated *GCN5* in transcription activation, and the encoded protein was subsequently shown to be the catalytic subunit of a HAT complex that primarily acetylates histone H3. SAGA’s 19 subunits are organized into submodules with distinct roles in histone acetylation (HAT module), histone deubiquitylation (DUB module), TBP binding (core module), and activator interactions (Soffers and Workman, 2020). In yeast, SAGA function is most important at inducible promoters, such as those activated by stress or certain nutritional conditions (Grant et al., 1997; Huisinga and Pugh, 2004). Chromatin immunoprecipitation and in vitro binding experiments show that transcription activators increase SAGA occupancy and histone H3 acetylation (H3ac) at promoters (Chen et al., 2012; Sikorski et al., 2012; Utley et al., 1998; Vignali et al., 2000). Bromodomains within the SAGA Spt7 and Gcn5 subunits recognize acetylated tails of both histone H3 and H4 (Li et al., 2007; Soffers and Workman, 2020), creating a positive-feedback loop and a possible activator-independent mechanism for directing SAGA to promoters. Importantly, while H3ac is strongest at the +1 nucleosomes of active genes, appreciable acetylation also occurs in downstream transcribed regions, leading to suggestions that SAGA also directly associates with transcription elongation complexes (Govind et al., 2007).

Co-localization Single Molecule Spectroscopy (CoSMoS) images individual protein-nucleic acid interactions using multi-wavelength total internal reflection fluorescence (TIRF) microscopy (Friedman et al., 2006; Friedman and Gelles, 2015). Our CoSMoS experiments reveal at least two modes of SAGA association with templates. SAGA readily binds to naked DNA independently of activator, but with short dwell times. In contrast, SAGA association with chromatin templates depends on activator. Activator increases both SAGA association rates and dwell times on both types of templates. SAGA dynamics are strongly affected by the specific activation domain, most consistent with direct, sustained interaction between the two factors, and arguing against the hit-and-run model. These single-molecule results thus provide a kinetic insight into how activators “recruit” the SAGA coactivator.

## RESULTS

### Single-molecule imaging of transcription activator and SAGA

Three color single-molecule microscopy was used to study activator-mediated recruitment of SAGA. The blue-excited channel was used to map locations of the DNA templates bound to the slide surface, red-excited for the transcription activator Gal4-SNAP-VP16, and green-excited for the coactivator SAGA (**Figure 1A**). By mapping molecule positions and subtracting background binding to “off-target” areas of the slide with no DNA, colocalizations can be used to quantitatively measure association and dissociation dynamics (Friedman and Gelles, 2015). To fluorescently label SAGA, a yeast strain was constructed (**Table S1**) in which the SNAP_f_ tag (Keppler et al., 2003; Keppler et al., 2004) was genetically fused to the C-terminus of SAGA subunit Spt7 (Baek et al., 2022). Endogenously expressed Spt7-SNAP was covalently labeled with SNAP-surface 549 dye (DY549, visible in the “green” channel) during nuclear extract preparation as previously described (Baek et al., 2021; Rosen et al., 2020). Yeast strains containing Spt7-SNAP showed no growth defects (**Figure S1A**). Nuclear extract from this strain showed specific dye labeling solely on Spt7 (**Figure S1B**) and was active for in vitro transcription (**Figure S1C**). To create a fluorescent transcription activator, a his_6_-tagged protein fusing the Gal4 DNA binding domain (DBD) to SNAP_f_ and the VP16 activation domain (hereafter Gal4-SNAP-VP16) was expressed in bacteria, purified, and labeled with SNAP-surface 649 dye (DY649, visible in the “red” channel). Insertion of the SNAP tag between Gal4 DBD and VP16 did not diminish transcription activation (**Figure S1C**, lanes 4-6).

**Figure 1.**
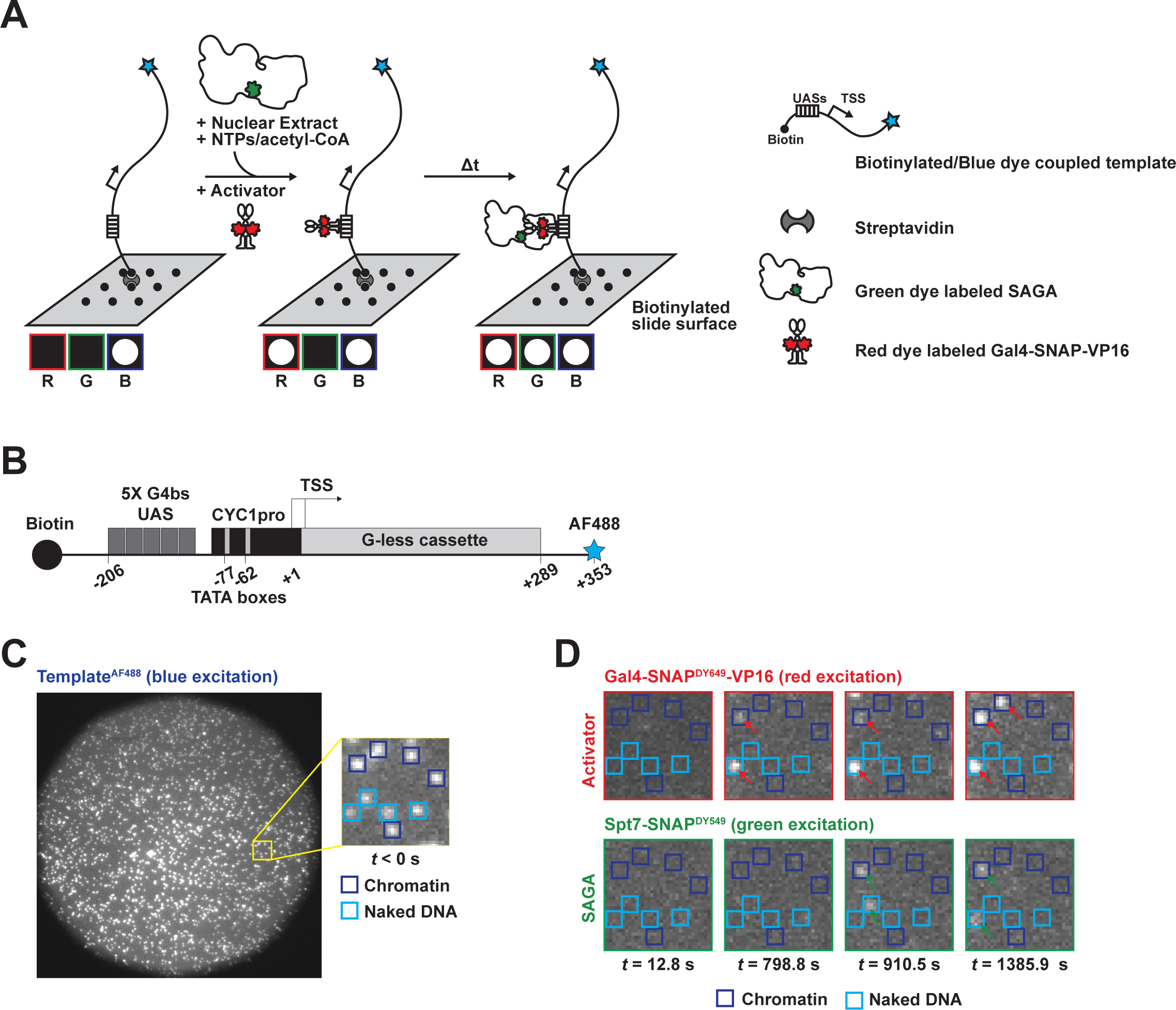
Single-molecule analysis of transcription factor association with the DNA template. **A.** Schematic diagram of CoSMoS system. DNA template molecules labeled with blue dye (AF488, blue star) are attached to the microscope slide via biotin-streptavidin-biotin linkage. After mapping the DNA locations, red-labeled transcription activator (Gal4-SNAP^DY649^-VP16) and nuclear extract containing green-labeled SAGA (Spt7-SNAP^DY549^) are flowed into the reaction chamber. Imaging the reaction with TIRF microscopy, colocalization of protein fluorescent spots with DNA locations are interpreted as binding events. **B.** Schematic of transcription template, with a UAS consisting of 5 Gal4 binding sites (G4bs) upstream of the CYC1 core promoter (CYC1pro) and a G-less cassette. Templates have biotin on one end for slide immobilization and blue dye AF488 on the other for visualization by TIRF microscopy. **C.** Microscope field of view showing DNA locations before incubation with proteins (t<0). Note that many experiments in this paper simultaneously assay binding to two DNA templates. One DNA (in this case, naked transcription template, boxed in light blue) is attached and imaged first to map its positions, followed by attachment and imaging of the second template (here, chromatinized transcription template). Yellow box shows area magnified in inset and panel D. **D.** Selected frames from time course. Top row (red) shows binding of activator, while bottom (green) shows SAGA binding. Numbers underneath (t) correspond to image time, and colored arrows indicate presence of bound protein.

The transcription template contains a UAS with five Gal4 binding sites and the *CYC1* core promoter upstream of a G-less transcription cassette (**Figure 1B**). Template DNA was PCR amplified using a 5’ biotinylated upstream primer and a 5’ Alexa Fluor 488 (AF488)-labeled downstream primer, tethered to a passivated microscope slide surface using a biotin-streptavidin-biotin sandwich, and visualized in the “blue” channel by laser excitation at 488 nm (**Figure 1C**). In some experiments, two different templates were compared on the same slide (e.g. naked versus chromatinized DNA, see below). For these experiments, the first template was attached and imaged, the slide chamber was extensively washed, and then the second template was immobilized and mapped in the same way. By this sequential image acquisition, naked (**Figure 1C**, cyan) and chromatinized (blue) templates were distinguishable in subsequent analyses. As “off-target” negative controls, hundreds of template-free slide surface areas were selected to measure non-specific surface binding by proteins of interest.

After mapping DNA positions, nuclear extract containing Spt7-SNAP^DY549^ and Gal4-SNAP^DY649^-VP16 was introduced into the microscope slide chamber (**Figure 1D**). Unless otherwise noted, reactions also contained 400 μM nucleotide triphosphates (NTPs) and 20 μM acetyl coenzyme A (acetyl-CoA) to mimic nuclear conditions. Over a time-course of roughly fifteen to twenty minutes (1000-1500 s), arrival and departure of Gal4-SNAP^DY649^-VP16 and Spt7-SNAP^DY549^ were monitored by alternating red (633 nm) and green (532 nm) excitation lasers with a cycle time of 1.4 seconds. Colocalization of proteins with DNA was determined using fluorescence intensity and proximity thresholds as previously described (Friedman and Gelles, 2015).

### SAGA dynamics on naked DNA

SAGA dynamics were first analyzed on naked DNA templates, comparing binding in the presence or absence of activator. Representative time records of Spt7-SNAP^DY549^ fluorescence intensity colocalizing with individual DNAs showed excellent signal-to-noise ratios (**Figure 2A**), and binding events of various durations were observed both in the presence (**upper panel, dark green**) or absence (**lower panel, light green**) of activator. There were also occasional periods when a second intensity jump (**Figure 2A**, black bars above trace) signified two molecules of SAGA bound to a single DNA molecule.

**Figure 2.**
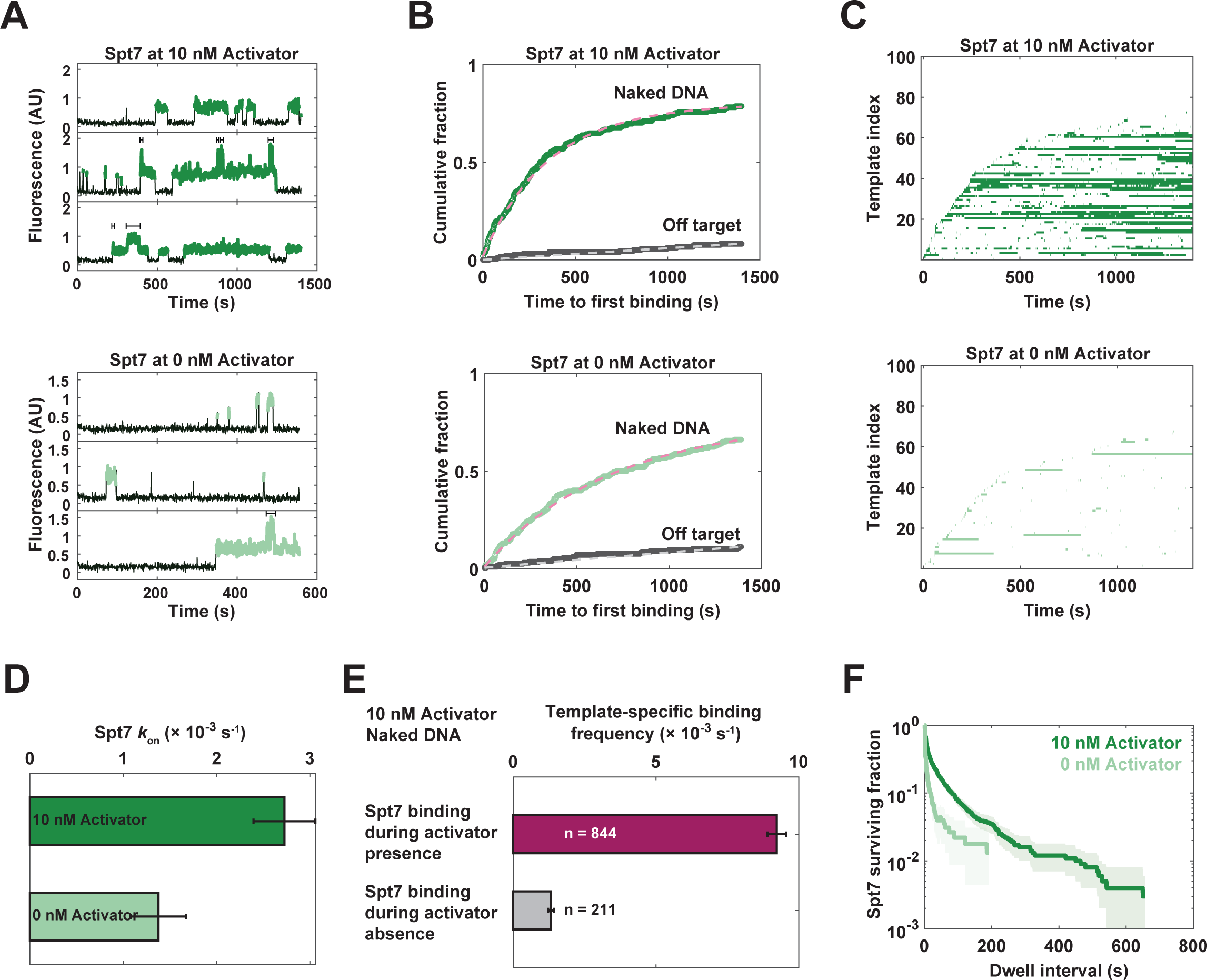
Activator-independent and activator-stimulated binding of SAGA to naked DNA templates. **A.** Intensity traces of Spt7-SNAP^DY549^ binding to naked DNA. Top panel shows binding at three individual DNAs in the presence of 10 nM Gal4-SNAP-VP16. Green color denotes presence of an Spt7-SNAP^DY549^ fluorescent spot, and brackets indicate multiple molecules present (seen at 46 out of 201 DNAs at this activator concentration). Bottom panel is the same, except in the absence of activator. **B**. Cumulative time-to-first-binding curves for SAGA, in the presence (dark green, upper panel) or absence (light green, lower panel) of activator. Non-specific background binding to slide locations lacking DNA is shown in gray. Note that these curves are derived from monitoring several hundred locations. Dashed lines show curve fits used for deriving the apparent association rates shown in panel D. **C.** Rastergrams showing SAGA binding to one hundred randomly chosen DNAs, ordered from bottom to top by time of first SAGA binding. Colored bars indicate intervals when one or more Spt7-SNAP^DY549^ molecules colocalized with the DNA molecule. **D.** SAGA association rates (± S.E.) calculated from time-to-first-binding curves in panel B. Error bars were calculated using bootstrapping. **E.** Frequencies of Spt7-SNAP^DY549^ arrivals to unbound DNA during intervals when activator was absent (gray) or present (purple). Error bars show standard error, and n indicates the number of arrivals used for each calculation. **F.** Survival plot for time intervals when one of more Spt7-SNAP^DY549^ molecules were bound to DNA. Data are corrected for non-specific surface binding by subtracting the off-target surviving fractions (determined by SAGA detection at spots with no mapped DNA). Note that a single exponential decay rate produces a straight line, so an inflection indicates two components representing species with distinct decay rates. Shaded areas represent the 90% confidence interval as determined by bootstrapping.

To better visualize the full set of binding events recorded, two representations of the data were used. In the first, the cumulative fraction of bound templates was plotted as a function of time to first SAGA binding at each DNA (**Figure 2B**). In the second, individual DNA binding records were converted to binary format, with color indicating when at least one Spt7 was present on the template. The resulting horizontal time ribbons were then stacked to form a rastergram, sorted (bottom to top) by the order of first Spt7-SNAP^DY549^ binding (100 randomly chosen DNAs are shown in **Figure 2C** and subsequent rastergrams, although several hundred DNAs are typically visualized in each experiment). Note that the leading edge of binding events created by the rastergram mirrors the cumulative time-to-first-binding curve, while the rastergram also conveys dwell time information.

To quantitate the effects of Gal4-SNAP^DY649^-VP16 activator, SAGA association rate constants were first derived by curve fitting to the cumulative time-to-first-binding plots (**Figure 2B**, dashed lines; note that all parameters for this and all subsequent curve fittings are listed in **Table S1**). This method avoids misinterpretation of any dye blinking during the experiment as a new binding event. Roughly 80% of DNA templates were bound by SAGA at least once in reactions containing 10 nM activator (**Figure 2B**, top panel, dark green curve). Unexpectedly, even in the absence of activator, ∼70% of templates colocalized with Spt7 at least once (bottom panel, light green curve). Spt7 binding fit very well to a single-exponential specific binding model (Friedman and Gelles, 2015), in the presence or absence of activator, suggesting a single rate-limiting step. At 10 nM activator, the apparent first-order association rate constant for Spt7 binding was 2.8 ± 0.3 × 10^-3^ s^-1^, whereas the rate without activator was 1.5 ± 0.3 × 10^-3^ s^-1^ (**Figure 2D**). Thus, using this time-to-first-binding method, the activator stimulates the initial association of Spt7 with naked templates only about two-fold under these conditions. Importantly, Spt7 showed very few binding events at off-target locations lacking DNA. Fewer than 10% of “off-target” spots showed any SAGA colocalization over the time course (**Figure 2B**, gray lines), while the majority of DNAs were bound at least once. We conclude that SAGA can specifically and efficiently bind to naked DNA template independently of activators.

As a second method for quantitating activator effects on SAGA, binding frequencies were calculated by dividing the number of all Spt7 arrivals by the combined time duration of the Spt7-free intervals (**Figure 2E**). In contrast to the method in **Figure 2B**, this approach incorporates both initial and subsequent Spt7 binding events. Note that neither method counts associations of second SAGA molecules to DNAs with one SAGA already bound. Because the activator was also labeled, Spt7 binding events could be separated by whether activator was present or absent on the DNA at the time of SAGA binding (see **Figure S1** for representative intensity traces). Using this method, activator produced a seven-fold increase in SAGA association rate, significantly higher than measured using only initial SAGA binding events. This quantitative difference occurs because, at 10 nM activator, a large fraction of initial Spt7 binding events occur before activator has bound the template (see below), and therefore do not reflect activator function.

In addition to the effects on association rates, the rastergrams show that activator produces a notable increase in average dwell time when one or more SAGA molecules are bound (compare bar lengths in **Figure 2C**). To quantitate this effect, the fraction of surviving binding events was plotted versus dwell time, after subtracting the off-target background values (**Figure 2F**). The inflected curves reveal two components that fit a bi-exponential decay: a larger fraction of bound molecules dissociate within a few seconds, while a smaller fraction persist on the order of a minute or more. Without activator (light green), ∼95% of SAGA binding events last less than 10 seconds. In the presence of 10 nM activator (dark green), about a third of Spt7 binding events are in the long duration fraction. While some of the extended durations result from overlapping occupancy by multiple SAGA molecules (see **Figure 2A**), it is clear that Gal4-SNAP-VP16 activator significantly increases average SAGA dwell times on the template.

### Co-imaging activator and SAGA on naked and chromatinized templates

Three color CoSMoS experiments were carried out using Spt7-SNAP^DY549^ extract, varying Gal4-SNAP^DY649^-VP16 activator concentrations from 0 to 30 nM. For this titration, both naked and chromatinized DNA templates were sequentially immobilized on the same microscope slide. Importantly, background binding of activator to the slide was negligible (**Figure S1B**). Rastergrams of activator binding to naked promoter DNA showed the expected increase in association rate with increasing concentration (**Figure 3A, upper red panels**). Template occupancy also increased, saturating at about 80% of DNAs. The fluorescence intensity traces showed that higher activator concentrations also increased the time when multiple activators were bound to the same template (**Figure S1A**).

**Figure 3.**
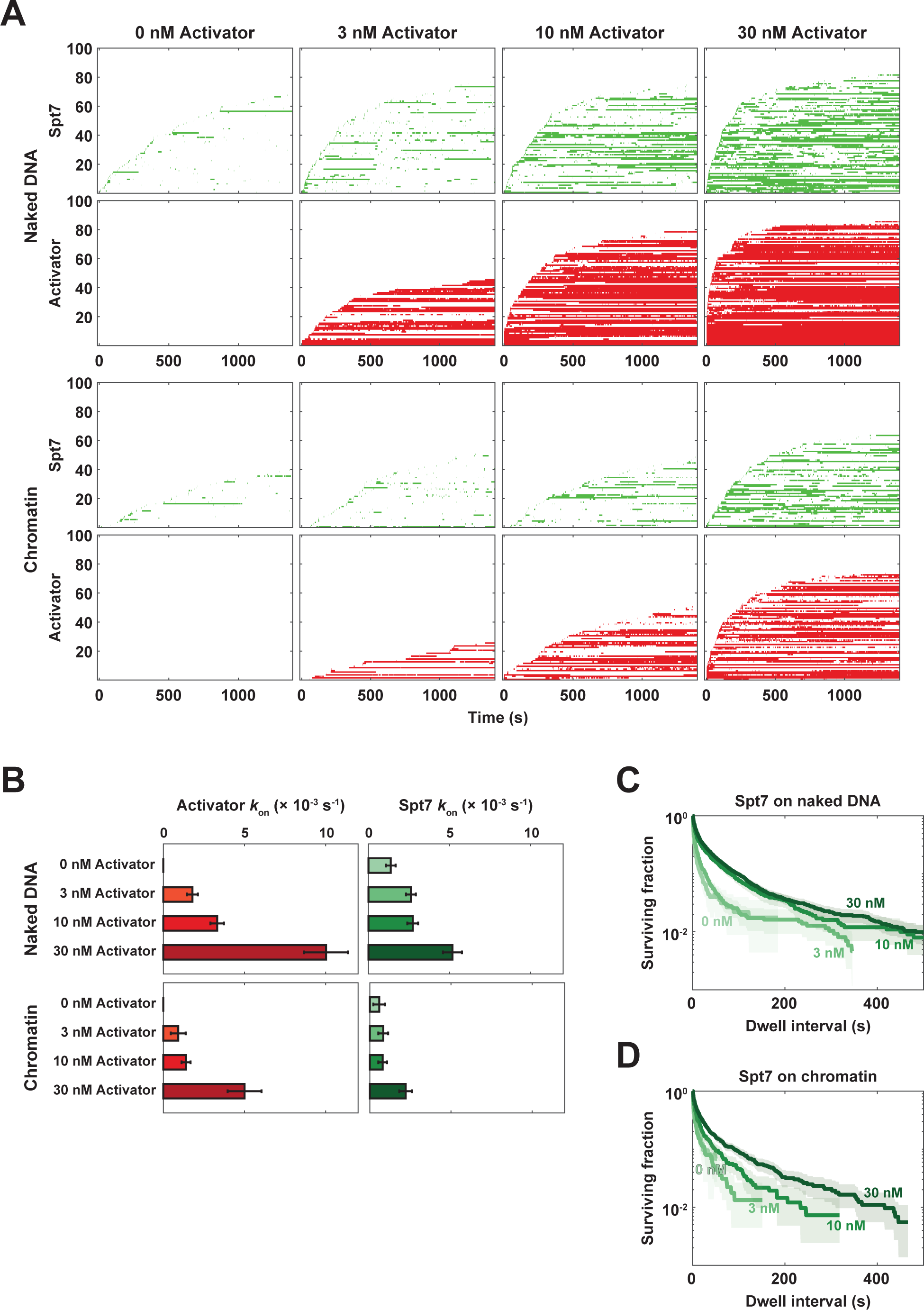
Effect of increasing activator concentration on SAGA binding to naked and chromatinized templates. **A.** Rastergrams of SAGA (Spt7, green) and Activator (red) binding. Naked and chromatinized templates were serially bound and mapped to the same slide surface, nuclear extract and the indicated concentration of activator were then flowed into the chamber, and imaging of the green and red channels initiated soon after (time zero). Y-axis numbers represent DNA index for 100 randomly chosen DNA or chromatin spots. **B**. Apparent association rates were calculated by curve-fitting a single exponential function to cumulative time-to-first-binding curves. Error bars showing standard deviation were determined by bootstrapping. **C**. Survival plot for SAGA dwell times on naked DNA at various activator concentrations (as in Figure 2F). Shaded areas show the 90% confidence interval as determined by bootstrapping. **D.** Same as panel C, except for SAGA binding on chromatinized DNA templates.

As before, Spt7-SNAP^DY549^ binding to roughly two-thirds of naked DNAs was seen in the absence of activator (**Figure 3A, first row, green**), far above background binding to off-target areas of the slide (**Figure S1B**). SAGA binding accelerated with increasing activator concentration. The association rate constants calculated by curve fitting to cumulative time-to-first-binding plots showed a non-linear response to activator concentration (**Figures 3B and S1C**). At lower activator concentrations of 3 and 10 nM, the initial rate of SAGA association increased no more than two-fold relative to no activator. However, at 30 nM Gal4-SNAP^DY649^-VP16, which nearly saturates the DNA at early time points, initial Spt7 binding was accelerated by roughly five-fold (**Figure 3B**). Because activator association at lower concentrations is slower than that of activator-independent SAGA binding, curve fitting to cumulative time-to-first-binding underestimates the activator response. We therefore calculated frequencies of total SAGA binding events to activator-bound versus unbound templates within each activator concentration condition. The presence of activator consistently led to 5- to 10-fold increase in the frequency of SAGA binding (**Figure S1D**).

In the absence of activator, dwell times when at least one SAGA molecule was present on naked DNA lasted on the order of just a few seconds (**Figure 3C**). As the concentration of activator was raised, the fraction of long-duration (∼1-3 minutes) Spt7 binding intervals markedly increased. However, the decay slopes of the activator-dependent long-duration events did not vary, indicating that the SAGA dissociation rate is independent of activator concentration. This result suggests that the same SAGA-activator-template interactions are made at all activator concentrations. In sum, these results show that activator both increases the rate of productive SAGA association with the template and slows its dissociation.

### Chromatin suppresses activator-independent SAGA binding

Nucleosomes inhibit interactions of many proteins with DNA, including those involved in transcription initiation and elongation (Cairns, 2009; Li et al., 2007). To examine the behavior of activators and SAGA in a more physiological context, the IswI-Nap1 system (Vary et al., 2004) and recombinant yeast histones were used to enzymatically reconstitute chromatin on the transcription template DNA (**Figure 4A**). Chromatin quality was confirmed by partial micrococcal nuclease digestion assays, where protected DNA fragments sizes indicated that 3-4 nucleosomes assembled on each DNA molecule (**Figure 4B**).

**Figure 4.**
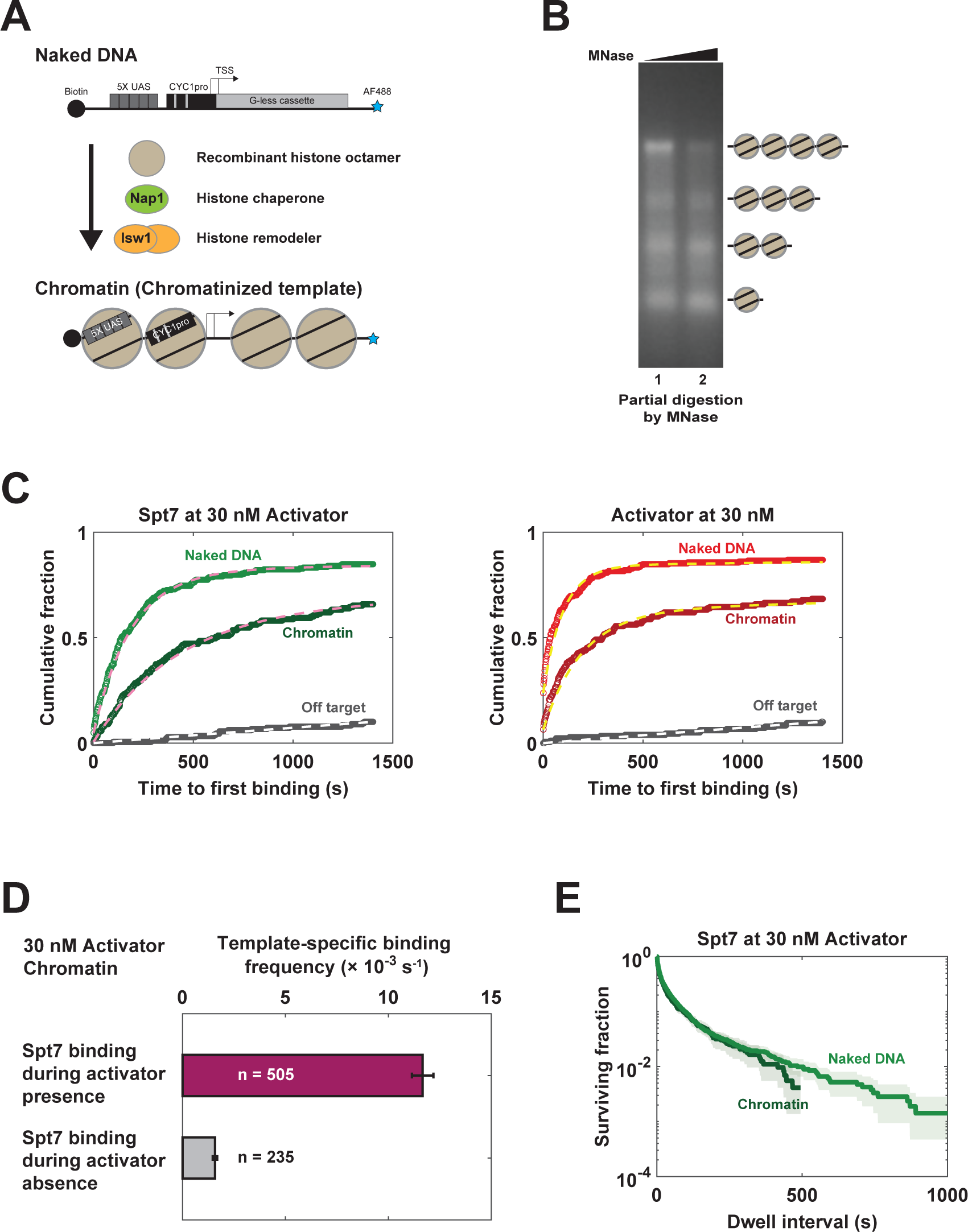
Analysis of SAGA binding to chromatinized DNA templates. **A.** Schematic of chromatin assembly process. Naked DNA template was incubated with histone chaperone Nap1, the chromatin remodeler Isw1a purified from yeast, and yeast histone octamers expressed in *E. coli*. **B.** Micrococcal nuclease digestion of chromatinized templates, followed by deproteinization and resolution of DNA by agarose gel electrophoresis. Limiting cleavage in linker regions reveals that each DNA carries up to four nucleosomes. **C.** Cumulative time-to-first-binding distributions for SAGA (Spt7-SNAP^DY549^, left panel) and Activator (Gal4-SNAP^DY649^-VP16, 30 nM, right panel) on naked (light colors) versus chromatin (dark colors) templates. Gray curves show off-target background binding to slide surface. Dashed lines show single exponential curve fits (parameters in Figure 3B and **Table S1**) **D.** Binding frequencies for SAGA binding on chromatinized templates when activator is present (purple) or absent (gray). Error bars show standard error, and n represents the number of events used in the calculation. **E.** SAGA survival fraction versus dwell time for naked and chromatin templates. Shaded areas indicate the 90% confidence interval as determined by bootstrapping.

With both chromatinized and naked DNAs immobilized on the same microscope slide, we could directly compare the binding behaviors for activator and SAGA (**Figure 3A**, upper versus lower rows). Consistent with earlier in vitro experiments (Taylor et al., 1991), Gal4-SNAP^DY649^-VP16 acts as a “pioneer” factor that can bind nucleosomal DNA, although its association rate was reduced by about half compared to naked DNA at all concentrations tested (**Figure 3B**). Some SAGA binding to chromatinized templates was again seen in the absence of activator, but greatly reduced compared to naked DNA. This binding might reflect incomplete chromatinization of some DNA molecules. SAGA binding on chromatin templates increased upon addition of 3 or 10 nM activator. Because of the slower activator binding, this increase was not strongly reflected in SAGA association rates derived from curve fitting to time-to-first-binding plots (**Figure 3B**), but calculation of the total SAGA binding frequencies showed a roughly 5-fold increase in the presence of activator, similar to the effect on naked DNA (**Figure S1D**). An apparent lag time was notable in the onset of SAGA binding to chromatin (**Figure 3A**), but not naked DNA. This lag interval decreased as activator concentration increased, suggesting that chromatin imposes a new rate-limiting step on SAGA binding that can be accelerated by activator (see below).

To best compare SAGA response to activators on naked versus chromatin templates, we used data from 30 nM activator conditions, where Gal4-SNAP^DY649^-VP16 rapidly binds both templates and SAGA binding fits a single exponential curve without a lag. Cumulative curves for first binding (**Figure 4C**) showed that both activator and SAGA association rates were decreased on chromatin, as were the total active fractions of templates bound (**Table S1**). However, on both types of templates, 30 nM activator produced a roughly 3-fold increase in SAGA association rate when calculated by curve fitting of initial bindings (**Figure 3B**), and a 6- to 8-fold increase in the frequency when measuring all SAGA binding events (**Figures S1D and 4D**).

In addition to accelerating SAGA association, activator also increased the proportion of long duration SAGA binding events on both naked and chromatin templates (**Figures 3C and D**). If long-lived binding events reflect only direct interaction of SAGA with the activator (see below), one would expect the dissociation curve slopes to be the same regardless of context. Indeed, at 30 nM activator, the dissociation slope for long-lived SAGA binding was the same on naked DNA or chromatin (**Figure 4E**, compare green and light green line slopes). However, SAGA dissociation was notably faster at lower activator concentrations on chromatin (**Figure 3D**), but not naked DNA (**Figure 3C**). What explains this difference? Dwell times when at least one Gal4-SNAP^DY649^-VP16 activator was bound also decreased on chromatin compared to naked DNA (**Figure S1**), consistent with earlier in vivo and in vitro studies on Gal4 suggesting nucleosome-induced dissociation (de Jonge et al., 2022; Donovan et al., 2019; Luo et al., 2014). Thus, the shorter duration of SAGA dwells on chromatinized DNA are likely indirectly due to activator dynamics.

The apparent lag in SAGA binding to chromatin at lower activator concentrations (**Figure 3A**) suggested an intervening rate-limiting step, most likely activator binding to nucleosomal DNA. To test whether activator preceded SAGA binding, two color rastergrams were created in which the first few seconds of initial activator binding on each DNA were marked in red, while the full duration of SAGA binding was marked in green (**Figure 5A**). While activator-stimulated SAGA binding was apparent on both naked and nucleosomal templates, chromatin clearly suppressed activator-independent SAGA binding (compare green SAGA binding events in the area above and below the first activator binding line in red). This effect was also apparent in individual intensity traces, where activator binding typically preceded initial SAGA binding to chromatin (**Figure 5B**). Histograms of the time intervals between arrivals for overlapping activator and SAGA binding events (**Figure 5C**, blue bars) showed a strong bias for activator binding first, far more than the distribution expected by chance (pink bars, showing a control simulation in which experimental binding records for the two proteins on different DNAs were randomly paired). Note that this bias is seen on both naked and chromatin templates, as expected from the strong activator-stimulated increase in SAGA association rates on both kinds of templates (**Figure S1D**).

**Figure 5.**
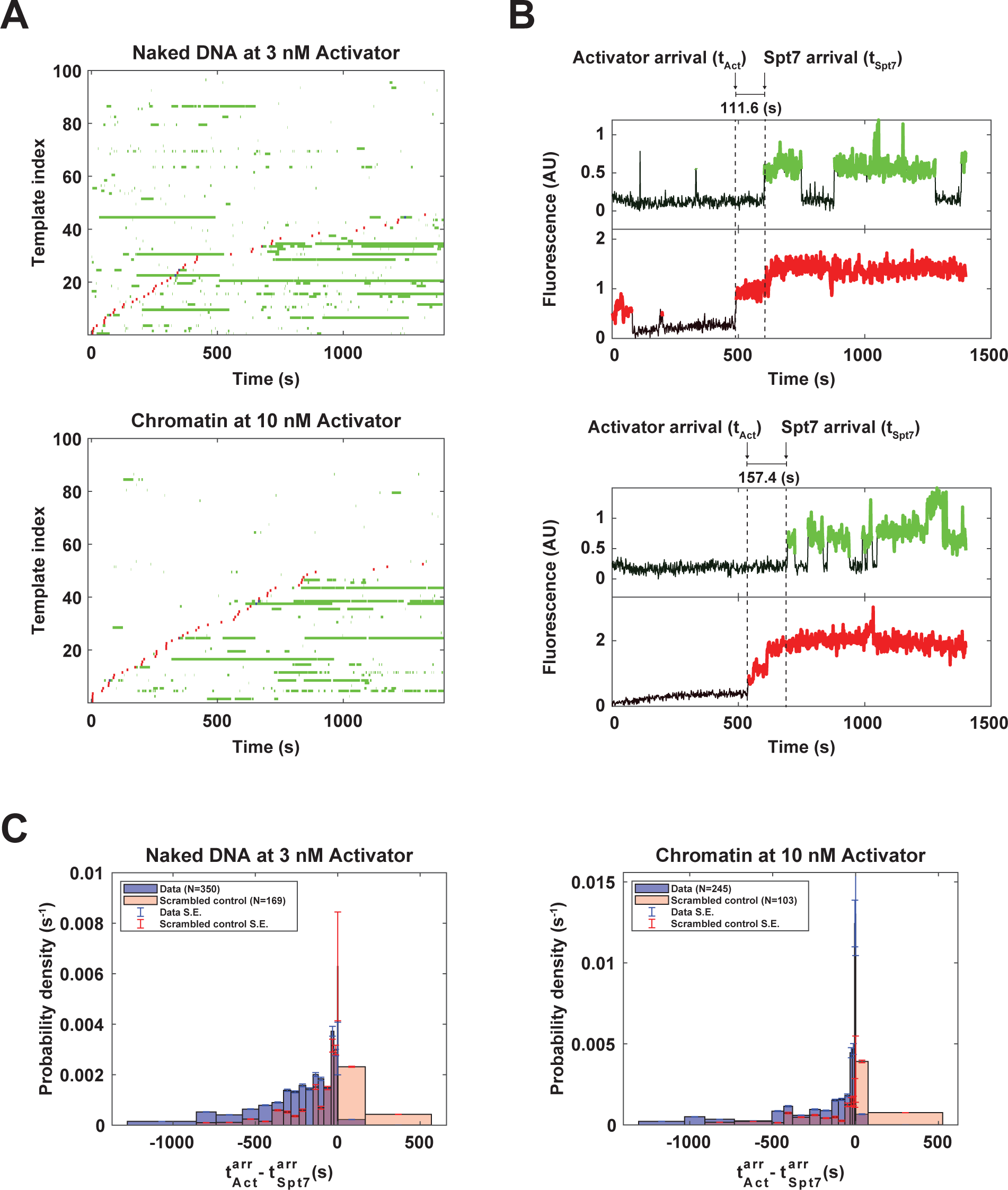
Chromatin suppresses activator-independent binding of SAGA to the template. **A.** Dual-color rastergrams show initial arrivals of Gal4-SNAP^DY649^-VP16 (red color marks first five frames of first activator binding on that DNA) and Spt7-SNAP^DY549^ (green shows entire SAGA-bound interval) to naked DNA templates (upper panel) or chromatinized templates (lower panel). Note that the activator concentrations for each type of DNA were chosen so that ∼50% of DNAs bound activator by the end of the experiment (see Fig. 3A). **B.** Intensity traces of two chromatinized templates showing how time intervals between SAGA (green) binding (t_Spt7_) and nearest activator (red) binding (t_Act_) were determined. **C.** Histogram showing the distribution of time intervals between overlapping SAGA and activator binding (t_Act_ - t_Spt7_). Blue shows actual data, while red is a “scrambled” control simulation where binding records for the two proteins on different DNAs were randomly paired. Error bars show standard error.

### Relative dwell times of activator and SAGA

In vivo tracking of single Gal4 molecules and other activators suggest brief residence on their binding sites, a few seconds at most (de Jonge et al., 2022). It is therefore pertinent to ask if the dwell times of individual activator molecules in our system are consistent with tethering SAGA for a minute or more. In one recruitment model, activator remains on the DNA, tethering SAGA for the entirety of the coactivator residence time. In an alternative “hit and run” model, activator stimulates the initial association of SAGA with the template, but is not required for maintenance, perhaps due to transfer of the coactivator to a nearby nucleosome or stretch of DNA. Combined rastergrams graphing periods of individual or overlapping binding show that long dwell times of Spt7-SNAP^DY549^ typically overlap periods when at least one Gal4-SNAP^DY649^-VP16 is bound (**Figure S1A**).

While consistent with a model where the coactivator remains tethered to the activator, this result does not rule out a hit-and-run model. One important caveat of our rastergram and dwell time analyses is they measure when at least one molecule of activator is bound to the five Gal4 binding site UAS. This means extended periods of activator occupancy might consist of multiple short duration binding events that overlap, and SAGA might transfer between these transient activator events without dissociating from the template. To measure dwell times of individual Gal4-SNAP^DY649^-VP16 molecules, a DNA template containing a single Gal4 binding site was used for CoSMoS. On naked DNA, in the context of nuclear extract, the 10 nM activator association rate was reduced by about half compared to the 5x Gal4 binding site template (compare left and middle panels in **Figure S1B** to the corresponding panels in **Figures 2B and 3A**, see **Table S1** for *k*_on_ rates calculated by fitting time-to-first binding curves), consistent with the decreased number of binding sites per template. Notably, both long and short duration bindings were observed, and these fit well to a biexponential dissociation curve (**Figure S1B,** right). Roughly 84% of activator dwell times were shorter than a few seconds. The remaining 16% lasted on the order of minutes, showing that a single activator can remain on the template long enough to tether SAGA for extended periods. The observation that the majority of SAGA and activator bindings last only a few seconds, but long duration SAGA binding is highly correlated with long duration activator binding, favors sustained tethering over a “hit-and-run” recruitment model.

### The transcription factor activation domain determines SAGA dynamics

Transcription activator function typically requires both a DNA binding domain (DBD) and a transcription activation domain. To validate that increased SAGA binding on the chromatinized template in the presence of Gal4-SNAP^DY649^-VP16 was not simply due to displacement of nucleosomes by the GAL4 DBD, SAGA binding was assayed in the presence of Gal4-SNAP^DY649^ lacking an activation domain. Binding dynamics of the two Gal4 derivatives were essentially identical (**Figure 6A**, bottom panels). In contrast, there was no stimulation of SAGA binding in the absence of the activation domain (**Figure 6A**, compare upper panels).

**Figure 6.**
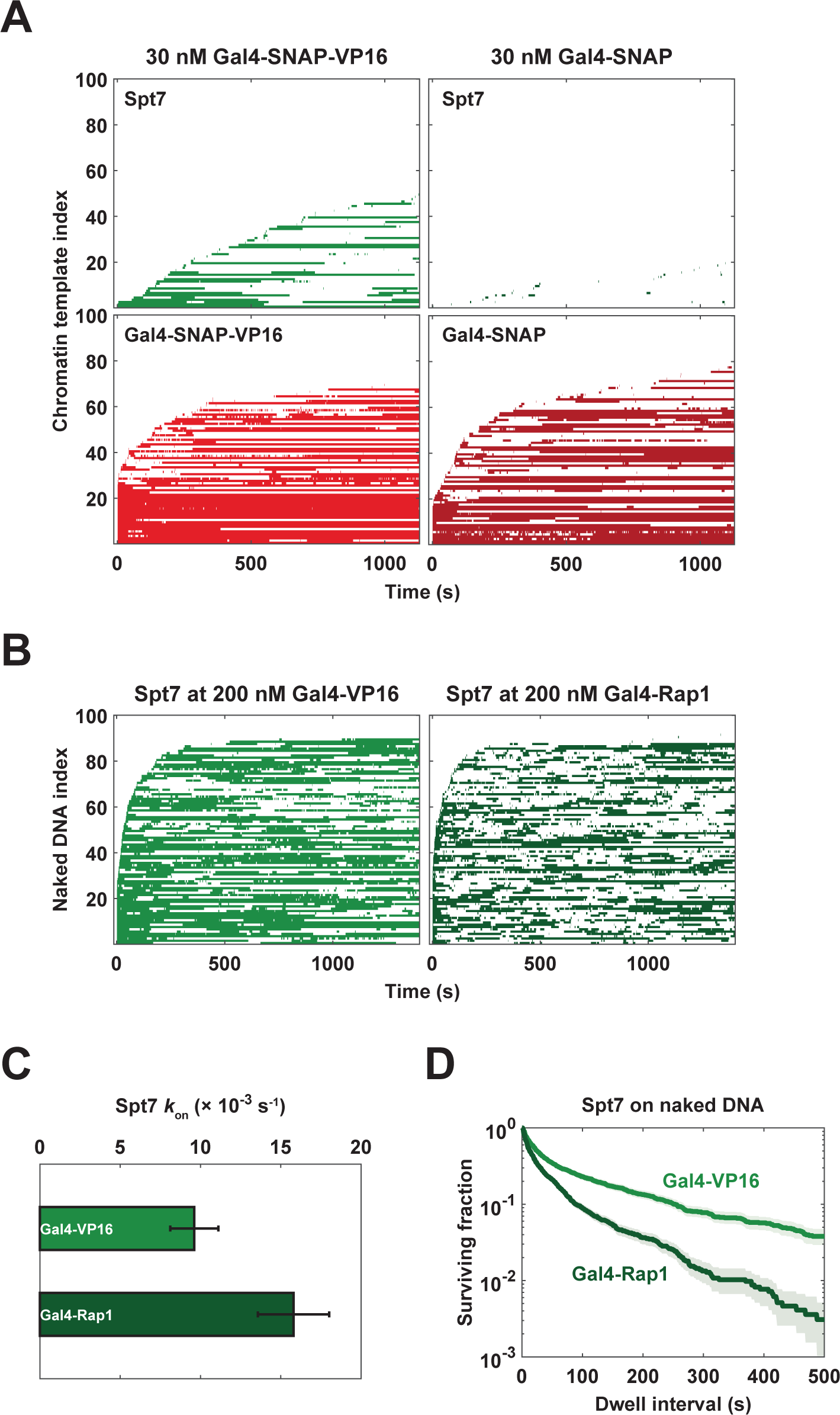
SAGA binding dynamics are determined by the activation domain. **A.** The Gal4 DNA binding domain is insufficient for SAGA recruitment. Rastergrams show binding data for SAGA (Spt7, green, upper panels) and the Gal4 derivative (red, lower panels), either when an activation domain was present (Gal4-SNAP^DY649^-VP16, left panels) or absent (Gal4-SNAP^DY649^, right). **B.** Rastergrams compare SAGA binding in the presence of high concentration (200 nM) Gal4 DNA binding domain derivatives fused to either the VP16 (light green, left) or Rap1 (dark green, right) activation domains, measured in the absence of NTPs. **C.** Association rates were calculated by curve fitting to the initial binding events of the experiment shown in panel B (see also **Figure S1A**). Error bars show standard error as determined by bootstrapping. Additional parameters are listed in **Table S1**. **D.** Dwell time survival curve of the full data set from the experiment in panel B was plotted. Shaded areas show the 90% confidence intervals as determined by bootstrapping.

Transcription activation domains are believed to directly contact coactivators. However, not all activation domains are equivalent. For example, we previously showed that the VP16 and Gcn4 activation domains are equally effective at recruiting SAGA and Swi/Snf to bead-immobilized DNA templates, but VP16 was much more effective at recruiting the NuA4 coactivator (Sikorski et al., 2012). To determine whether SAGA dynamics are sensitive to the particular activation domain used, we compared the Gal4 DBD fused to the activation domains from either VP16 or the yeast transcription regulator Rap1 (aa 630 – 695, (Hardy et al., 1992)). The activators were used at 200 nM to rapidly saturate template binding, and Spt7-HALO^JF646^ extract was imaged (rastergrams shown in **Figure 6B**). Interestingly, the SAGA association rate was significantly faster with Gal4-Rap1 AD (**Figures 6C and S1A**), but so was the apparent dissociation rate (**Figure 6D**). The sensitivity of SAGA dwell time to the particular activation domain rules out a model in which the activator indirectly recruits the coactivator by creating an altered state of the template (for example, a stretch of naked DNA) that binds SAGA. Instead, these results are most consistent with a model where activator directly tethers SAGA to the template through sustained contact.

### Effects of NTPs and acetylation on SAGA and activator binding

HATs and ATP-dependent chromatin remodelers are known to cooperate as transcription coactivators. To provide for these enzymatic activities, the experiments presented above contained 400 μM of each of the four NTPs and 20 μM acetyl-CoA to mimic in vivo conditions. To test the influence of histone acetylation on activator-SAGA dynamics, chromatin templates were assembled with semi-synthetic recombinant human histones. Unmodified nucleosomes were compared to those tetra-acetylated on either histone H4 (K5, K12, K16, K20, **Figures 7A-D, S1A+B**) or H3 (K4, K9, K14, K18, **Figures S1C+D**). Modified and unmodified templates were sequentially immobilized on the microscope slide and imaged in nuclear extract containing Spt7-SNAP^DY549^ and 20 nM Gal4-SNAP^DY649^-VP16.

**Figure 7.**
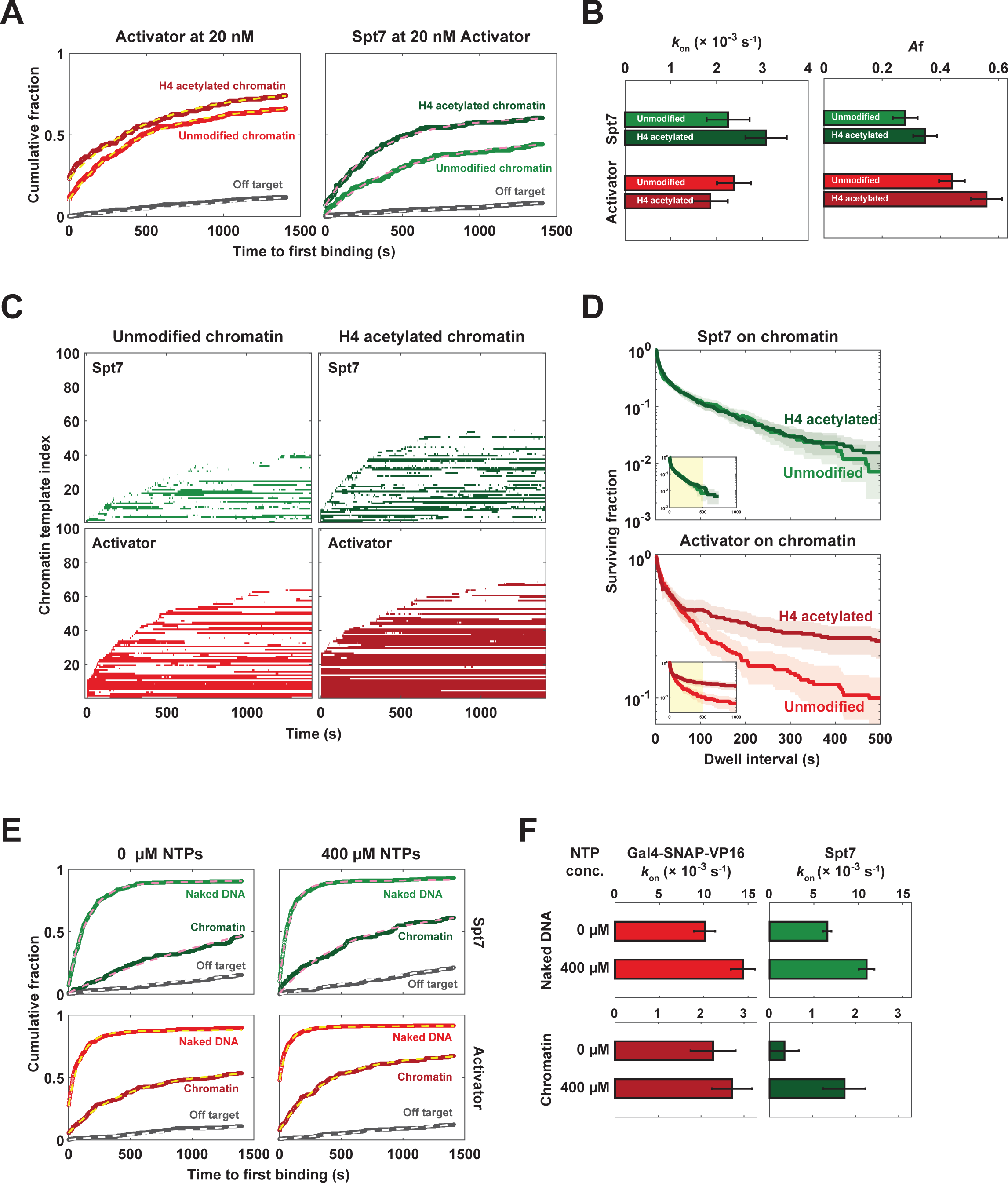
NTPs and histone acetylation affect both SAGA and activator binding. **A.** Cumulative time-to-first-binding curves are shown for activator (Gal4-SNAP^DY649^-VP16, red) and SAGA (Spt7-SNAP^DY549^, green) binding to templates assembled with unmodified (light colors) or H4 tetra-acetylated (dark colors) chromatin templates, or to slide areas without DNA (Off-target, gray). The two templates were sequentially immobilized on the same microscope slide for direct comparison. **B.** Active template fraction (right panel) and association rate parameters (left) were derived by curve fitting to data in panel A (see dashed lines). See also **Table S1**. Bars show standard error as determined by bootstrapping. **C.** Rastergrams show factor occupancy intervals from the same experiment as panel A. **D.** Dwell time survival curves for the first 500 seconds of data from panel are shown, with the first 1000 seconds of data shown in the inset. Shaded areas with green and red colors show the 90% confidence intervals. **E.** Cumulative time-to-first-binding curves are shown for activator (Gal4-SNAP^DY649^-VP16, red, bottom panels) and SAGA (Spt7-SNAP^DY549^, green, upper panels) to naked DNA templates (light colors), chromatin DNA templates (dark colors), or slide areas without DNA (Off-target, gray). Left panels show results in the absence of NTPs and right panels in the presence of 400 μM NTPs. **F.** Association rate parameters were derived by curve fitting to data in panel E (see dashed lines). See **Figure S1F** for rastergrams and **Table S1** for additional parameters. Bars show standard error as determined by bootstrapping.

Given SAGA’s two bromodomains, we expected increased dwell times on the acetylated chromatin. In the absence of added NTPs and acetyl-CoA, SAGA and activator binding were unaffected by H4 acetylation (**Figures S1A+B**). However, in the presence of NTPs, SAGA association increased on H4ac relative to unmodified templates (**Figure 7A, right panel, Table S1**), while dissociation was unaffected (**Figure 7D, upper panel**). In contrast, activator association was unaffected, but the duration of intervals when at least one activator occupied the template was unexpectedly significantly increased (**Figure 7D, lower right panel**). The H3ac templates also showed increases in duration of activator-bound periods and SAGA association, although these effects did not require NTPs (**Figures S1C+D**).

We interpret these unexpected results as follows. As a pioneer factor, initial binding of Gal4-SNAP^DY649^-VP16 may be insensitive to histone acetylation. However, once bound, the activator can recruit Swi/Snf or possibly other ATP-dependent chromatin remodelers. The acetylated histone tails might further promote remodeler binding through interactions with the bromodomains found in Swi/Snf and RSC. NTPs could then trigger nucleosome movement or displacement in a manner that prolongs activator occupancy, either by extending the dwell time of individual Gal4-SNAP^DY649^-VP16 molecules, or more likely, by exposing additional Gal4 binding sites so multiple activators can bind. The increased activator occupancy would in turn explain the increased SAGA association. As for dissociation, there may be a slight stabilization of SAGA on H3ac chromatin (**Figure S1D**), but H4ac had no effect (**Figure 7D**). These results are consistent with the fact that SAGA bromodomain mutants have only minor effects in vivo (Sterner et al., 1999), and together with our earlier experiments, argue that longer duration SAGA binding is primarily mediated by contacts with the activator rather than the chromatin itself.

This model predicts that the NTP stimulation should be specific to chromatin templates. Indeed, direct comparison of naked and unmodified chromatin templates showed that SAGA association kinetics, calculated either by initial binding curve fitting (**Figures 7E+F, S1F**), or total binding frequency (**Figure S1E**), showed a much stronger response to NTPs on chromatin. Note that this experiment included acetyl-CoA to support histone acetylation. The observation that NTPs have less stimulatory effect on naked DNA, for both SAGA and activator binding, is consistent with template-associated histones being displaced by remodelers or other NTP-dependent mechanisms. Further supporting our model, the stimulatory effect of NTPs on SAGA binding to chromatin templates is only seen when activator is bound (**Figure S1E,** bottom panel). Future experiments will be needed to determine whether the observed effects require all NTPs or only ATP, and whether they reflect nucleosome remodeling, transcription, or other NTP-dependent functions. No matter the mechanism, our results illustrate that both activator and co-activator binding are dynamic processes unlikely to be accurately represented by simple equilibrium binding assays with purified proteins.

## DISCUSSION

Our single-molecule results provide kinetic insight into how sequence-specific transcription factors “recruit” the co-activator SAGA. The data reveal two modes of association of SAGA with transcription templates. SAGA readily associates with naked DNA in the absence of activator, but with a relatively short dwell time (**Figures 2 and 3**). This “basal” binding mode is inhibited by nucleosomes, but as discussed below, is likely to be physiologically relevant. In the alternative “activator-tethered” mode, interaction of SAGA with a transcription activation domain both increases the SAGA association rate and increases SAGA dwell time (**Figures 2-4**). We did not find strong evidence for a proposed third mode of binding in which the SAGA is tethered to chromatin via bromodomain interactions with acetylated histone H3 and H4 tails, independently of activator (**Figures 7A-D**). With a clearer understanding of factor dynamics, better models of transcription activation now emerge.

### SAGA interactions with nucleosome-free DNA

Activator-independent binding of SAGA to naked DNA might be dismissed as an artifact, but several facts argue against this. First, a quantitative analysis of in vitro acetylation kinetics found that SAGA activity on mono-nucleosomes is strongly accelerated (5- to 9-fold) by unwrapped DNA flanking the target nucleosome (Mittal et al., 2018). Importantly, the minimal stimulatory DNA length exceeds the typical in vivo inter-nucleosomal linker size. Second, ChIP-exo experiments from the Pugh lab suggest that many yeast promoters function without detectable sequence-specific regulatory transcription factors or co-activators (the UNB class in Rossi et al., 2021). Although this conclusion is controversial, most yeast promoters do reside within intrinsically nucleosome-free or -depleted regions. These stretches of unwrapped DNA may directly bind SAGA, albeit with short dwell times that limit detection by crosslinking, thus explaining how promoters with little cross-linkable SAGA still have significant H3 acetylation on flanking nucleosomes. Finally, ChIP experiments reveal lower but detectable levels of both SAGA and H3 acetylation within coding regions of genes, far from activator-binding sites. It is proposed that direct contacts between SAGA and RNApII mediate promoter-distal acetylation (Govind et al., 2007), but mass spectrometry analysis of elongation complexes did not detect SAGA (Joo et al., 2019a). Our results suggest transiently exposed DNA resulting from transcription could directly recruit SAGA to modify an adjacent nucleosome. We suspect that DNA exposure by transcription or chromatin remodeling may underlie the observed stimulation of SAGA binding by NTPs/ATP (**Figures 7E, 7F, S1E, 7F**).

### SAGA tethering to transcription activators

Although it is well-established that transcription activators interact directly with SAGA to “recruit” the HAT to promoters, the experiments underlying this conclusion typically measure binding in aggregate under equilibrium conditions. Our single-molecule experiments reveal that activator-mediated “recruitment” of SAGA has two components: an increase in association rate, and a significant decrease in dissociation rates (**Figures 2E-F, 3B-D, S1D**). Both of these effects are mediated by direct interactions with the transcription factor activation domain. The Gal4 DNA binding domain alone produces no stimulation (**Figure 6A**), ruling out a model where activator binding to DNA simply displaces histones to generate naked DNA. More compellingly, notable differences in association and dissociation rates are seen between the VP16 and Rap1 activation domains (**Figures 6B-D**). These differences suggest the activation domain does not just deliver SAGA in a “hit and run” type mechanism, but instead retains contact with SAGA while the coactivator is bound. It is thus possible that SAGA occupancy can be differentially tuned at individual promoters by which activators are bound at a given time.

Although the activator-bound and DNA-bound states could theoretically be mutually exclusive, SAGA may be capable of simultaneous interactions because the interaction surfaces appear non-overlapping. The Tra1 subunit interacts with activation domains (Hahn and Young, 2011), while the SWIRM domain of Ada2 has been proposed to directly bind DNA (Qian et al., 2005). Consistent with this dual-interaction model, Mittal et al (Mittal et al., 2018) used an in vitro mono-nucleosome assay to show that binding of a single Gal4-VP16 produced a five-fold increase in histone acetylation, on top of a four-fold increase already seen upon addition of 80 bp of unwrapped flanking DNA. Interestingly, that study concluded that activator did not change SAGA affinity for the nucleosome (as measured by the *K*_m_ for acetylation), but instead stimulated the catalytic rate (*k*_cat_). In apparent contradiction, our results show clear activator effects on SAGA affinity. However, we measured binding to the transcription template as a whole, rather than the nucleosome per se, and in the very different context of nuclear extract on templates having multiple activator binding sites and nucleosomes.

### Models for targeting histone acetylation

Given the two modes for SAGA interaction with the transcription template, three models arise for how acetylation may be targeted by activators (**Figure S1B**). In model 1, activator-bound SAGA is an intermediate from which the HAT is transferred to nearby exposed DNA. Acetylation of flanking nucleosomes would then be performed by DNA-bound SAGA. By creating a high local concentration of coactivator near the enhancer/UAS, activators would facilitate delivery of SAGA to nearby promoters or other regions of naked DNA where nucleosomes have been displaced by remodelers or transcription. In the second model, SAGA acetylates nucleosomes while remaining bound to the activator, using the flexibility of the activation domain and the DNA itself to reach its substrate. Here, SAGA contacts only the activator and the substrate histone, with no DNA binding required. In the third model, activator-tethered SAGA simultaneously contacts naked DNA while acetylating the target nucleosome.

Because any model for acetylation targeting must explain how coactivators bound at distant enhancers acetylated their target promoters but not all the intervening chromatin. SAGA’s affinity for naked DNA, together with the strong stimulation of SAGA HAT activity by DNA flanking the substrate nucleosome (Mittal et al., 2018), suggests a simple targeting mechanism. We therefore favor models 1 and 3, in which acetylation occurs while SAGA interacts with naked DNA, while either tethered to, or recently released from, the activator. CoSMoS only detects binding, so future experiments distinguishing these two models will require an approach that reports on both activator interaction and the exact timing of acetylation.

### Dynamics of activators and coactivators on chromatin

On a single Gal4 binding site template, the majority of activator binding events last only a few seconds, while the remaining 10-20% dissociate on the order of minutes (**Figure S1B**). Single molecule tracking (SMT) of Gal4 and other transcription activators in vivo typically detects three behaviors: free diffusion, ∼1 second pauses that are thought to reflect non-specific DNA interactions, and pauses on the order of tens of seconds believed to represent sequence-specific binding (de Jonge et al., 2022; Donovan et al., 2019). The majority of Gal4 dwell times seen by CoSMoS are also in this range, but about 15% of binding events lasted on the order of minutes (**Figure S1B**). Notably, dye photobleaching and other technical issues makes it difficult to continuously track single molecules in vivo for more than a minute using SMT (de Jonge et al., 2022; Garcia et al., 2021). Recently, an improved tracking system showed that a previously undetected fraction of Gal4 binds the *GAL10* promoter in vivo with an average dwell time of 83 seconds (Pomp et al., 2023). Therefore, activator behavior in our system aligns well with in vivo results.

If each SAGA molecule is tethered to a single activator molecule, the SAGA dissociation rate should be independent of activator concentration. This appears to be the case on naked templates, where normalized SAGA survival curves for the 10 and 30 nM activator conditions are nearly identical, and the 3 nM sample, where most templates don’t have bound activator, resembles the 0 nM activator curve (**Figure 3C**). In contrast, the survival plots for SAGA on chromatin show different slopes for 10 versus 30 nM Gal4-VP16, indicating faster dissociation of SAGA at the lower activator concentration (**Figure 3D**).

One possible explanation for this unexpected behavior is that higher activator concentrations produces more periods where multiple activators, and therefore multiple SAGAs, are bound simultaneously. As the typical CoSMoS analysis only registers when at least one complex is bound, temporal overlap of multiple SAGA molecules can produce extended periods of binding. Alternatively, the dwell time for a single SAGA molecule might be extended if it can interact with multiple activators, either simultaneously or sequentially. Activation domains have both relatively low affinity and high promiscuity for targets, so SAGA may more than one activator interaction site. Alternatively, the close proximity of multiple Gal4-VP16 molecules on the UAS might facilitate rapid reassociation of any SAGA complexes that dissociate. However, for all of these models it is unclear why naked DNA templates, where multiple activator occupancies are just as frequent as on chromatin (**Figure S1A**), would not show the same effect.

A more likely explanation is that chromatin reduces the dwell time of activator molecules, which indirectly results in reduced SAGA occupancy time. It is well documented that nucleosomes accelerate Gal4 dissociation in vivo and in vitro (de Jonge et al., 2022; Donovan et al., 2019; Luo et al., 2014), and our system shows a similar effect for Gal4-SNAP-VP16 (**Figure S1**). Notably, higher activator concentrations abrogate this effect, as the SAGA dissociation rate is the same on naked and chromatin templates when 30 nM activator or higher is used (**Figures 3C and D**). Experiments comparing unmodified and acetylated nucleosomes provide another indication that activator residence time is highly sensitive to the chromatin environment. Surprisingly, despite its bromodomains, SAGA dissociation was unaffected. In contrast, activator occupancy times were significantly extended (**Figures 7D, S1B, D**), which likely accounts for the slight increase in SAGA association rates (**Figures 7A, B, S1A, C**). Deciphering the mechanism by which histone acetylation stabilizes activator binding will require future experiments, but we suspect it may involve one or more chromatin remodelers, as NTPs also contribute (**Figures 7E, F, S1E, F**). This model is consistent with the concept of a “pioneer” factor, whose initial binding can open up the adjacent chromatin for additional activator binding (Zaret, 2018).

### Future directions

We previously showed that RNApII and a subset of basal transcription factors pre-assemble on the UAS to form a pre-PIC for transfer to the core promoter (Baek et al., 2021). These complexes are tethered to the template via the activator-Mediator interaction (Gorbea Colon et al., 2023)(**Zhou/Jeon, in preparation**). It is remarkable that VP16 and many other eukaryotic activation domains can recruit not only Mediator, but also SAGA, NuA4, and Swi/Snf remodeler. Using our extract-based system, future CoSMoS experiments will test if these co-activators are recruited in a particular order, and whether any bind cooperatively to the activators at the UAS. The different co-activators must somehow contend with each other during the process of transcription activation, so it is important to understand their dynamics.

## METHODS

### Yeast strains

*S. cerevisiae* strains used in this study are listed in Table S1. Spt7-3HA-SNAP_f_ and Rpb1-3HA-HALO C-terminal tagging cassettes were PCR amplified with the appropriate plasmids (Table S1, (Baek et al., 2022)) and primer pairs (Table S1). The amplified DNA fragments were transformed into the protease-deficient strain, YF702/CB012 (Inada et al., 2002), or its derivatives already containing another fusion protein (YSB3337 and YSB3573), as indicated. Positive transformants selected by the insertion markers were further confirmed for proper insertion by colony PCR with the appropriate primer pairs from Table S1. SNAP_f_ and HALO fusion protein expression and stability were verified by immunoblotting. The fusion strains had similar growth to the YF702 wild type as determined by spotting assay (Figure S1A).

### Yeast nuclear extract preparation

Yeast nuclear extracts were prepared as previously described with slight modifications (Baek et al., 2021). Yeast cells were grown in 4 liters of modified YPD medium (1% yeast extract, 2 % peptone, 3% dextrose, 0.015% tryptophan, and 0.006% adenine) at 30°C to an optical density at 600 nm of 3–5 and were harvested by centrifugation at 4000 rpm (Sorvall GS3 or SLA-3000, Thermo Scientific) for 8 min. Cells were suspended in 150 ml of TD buffer (50 mM Tris pH 7.5, and 30 mM DTT (dithiothreitol)) and incubated with slow shaking at 30°C for 15 min. Cells were harvested by centrifugation at 4000 rpm for 12 min and resuspended in 20 ml of YPD containing 1M sorbitol. To digest cell walls, the resuspended cells were mixed with 15 mg of Zymolyase 100T (Seikagaku Corp.) that was dissolved in 30 ml 1 M sorbitol. After 30-60 minutes digestion at 30°C, when 80∼95% of cells became spheroplasts, digestion was stopped by addition of 100 ml of YPD containing 1 M sorbitol and centrifugation at 4000 rpm for 12 min. Spheroplasts were resuspended in 250 ml of YPD containing 1 M sorbitol, and incubated at 30°C with slow shaking for 30 min recovery time. Spheroplasts were next washed twice with cold YPD containing 1 M sorbitol and then with cold 1 M sorbitol, each time centrifuging at 4000 rpm for 12 min. Finally, spheroplasts were suspended in 100 ml Buffer A (18% (w/v) Ficoll 400, 10 mM Tris-HCl pH 7.5, 20 mM potassium acetate, 5 mM magnesium acetate, 1 mM EDTA (ethylenediamine tetraacetic acid) pH 8.0, 0.5 mM spermidine, 0.17 mM spermine, 3 mM DTT, and 1× protease inhibitors (1 μg/ml each of aprotinin, leupeptin, pepstatin A, and antipain)). The suspended spheroplasts were lysed with a motorized Dounce homogenizer (Wheaton, #62400-802). Cell debris and incompletely lysed spheroplasts were removed by four sequential centrifugations in a Sorvall GSA rotor (twice at 5000 rpm for 8 min followed by twice at 5000 rpm for 5 min) of the supernatant. Crude nuclei were pelleted by centrifugation at 13000 rpm in a Sorvall GSA rotor for 30 min, and then suspended in 10 ml of Buffer B (100 mM Tris-acetate pH 7.9, 50 mM potassium acetate, 10 mM magnesium sulfate, 10% glycerol, 2 mM EDTA pH 8.0, 3 mM DTT, and 1× protease inhibitors). The nuclei were lysed by the dropwise addition of 3 M ammonium sulfate solution (pH 7.5) to a final concentration of 0.5 M, followed by additional rotating at 4°C for 30 min. The lysed nuclei were centrifuged in a Beckman type 70 Ti rotor at 37,000 rpm for 1.5 hr at 4°C. The supernatant was collected, and the nuclear proteins were precipitated by the slow addition of granular ammonium sulfate to ∼75 % saturation (0.35 g to 1 ml of the supernatant). The suspension was incubated at 4°C for 30 min, and nuclear proteins were pelleted by centrifugation in a Sorvall GSA rotor at 13,000 rpm for 20 min, the supernatant was removed, and then the pellet was spun again for 5 min to collect at the bottom of the tube. The protein pellet (∼0.8 g) was suspended in 2 ml of Buffer C’ (20 mM HEPES pH 7.6, 10 mM magnesium sulfate, 1 mM EGTA (ethylene glycol-bis(β-aminoethylether) tetra-acetic acid), 10% glycerol, 3 mM DTT, and 1× protease inhibitors). For labeling of SNAP_f_- or HALO-fused Spt7, the nuclear extract was incubated with 0.4 μM SNAP-surface 549 (New England BioLabs, #S9112S) or HALO-JF646 (Luke Lavis, Jenelia Farms) at 4°C for 1 hr on a rotator in the dark. The labeled nuclear extract was then dialyzed against 500 ml of nuclear extract dialysis buffer (20 mM HEPES-KOH pH 7.6, 75 mM ammonium sulfate, 10 mM magnesium sulfate, 1 mM EGTA, 10% glycerol, 3 mM DTT, and 1× protease inhibitors) in a dialysis membrane (molecular weight cut-off, 6-8 kDa) three times (1 hr, 1.5 hr and 2 hr). Residual unreacted dye in the nuclear extract was removed by incubation at 4°C for 1 hr on a rotator with one fourth extract volume of agarose beads (Pierce 26196 or Thermo 20381) that were coupled to ∼10 mg SNAP or HALO protein, after which the beads were removed by table-top centrifugation at 1,000 x *g* for 2 min at 4°C (Baek et al., 2021; Haraszti and Braun, 2020). Nuclear extract aliquots were flash frozen with liquid nitrogen and stored at -80°C until use. Labeling of Spt7 and residual dye depletion were confirmed by in-gel fluorescence imaging on a Typhoon imager (GE Healthcare). *In vitro* transcription activity of the nuclear extract was verified by the *in vitro* transcription assay described below.

### Preparation of recombinant transcription activators

pRJR-Gal4-SNAP_f_-VP16 (BE559) and pRJR-Gal4-SNAP_f_ (BE562) plasmids were cloned by isothermal DNA assembly of PCR fragments amplified using the indicated parent plasmid templates (Table S1) and primers (Table S1), respectively. These fusions, as well as Gal4-SNAP_f_-VP16, Gal4-SNAP_f_-Rap1, and Gal4-SNAP_f_ were purified as previously described with slight modifications (Sikorski et al., 2012). BL21(Codon+, DE3) containing the appropriate expression plasmid was grown at 37°C in LB medium with appropriate antibiotics to an optical density at 600 nm of 0.5. Proteins were induced with 0.1 mM IPTG and 10 μM zinc chloride for 3 hr at room temperature. Bacteria were harvested by centrifugation at 5000 rpm (Sorvall GS3 or SLA-3000, Thermo Scientific) for 10 min at 4°C, resuspended in lysis buffer (20 mM HEPES–KOH pH 7.6, 300 mM potassium chloride, 10 μM zinc chloride, 5% glycerol, 0.1% NP-40, 1 mM PMSF), and lysed by sonication. The soluble extract was isolated after centrifugation at 15000 rpm (Sorvall GSA rotor) for 15 min at 4°C, and then incubated with Ni^2+^-NTA-agarose resin (Goldbio) overnight on a rotator at 4°C. The protein-bound resin was extensively washed with Wash Buffer (20 mM HEPES–KOH pH 7.6, 30 mM potassium chloride, 10 μM zinc chloride, 15 mM imidazole, 5% glycerol, 0.1% NP-40, 1 mM PMSF), and bound proteins were eluted with Elution Buffer (20 mM HEPES–KOH pH 7.6, 30 mM potassium chloride, 10 μM zinc chloride, 600 mM imidazole, 5% glycerol, 0.1% NP-40, 1 mM PMSF). The supernatant was clarified by centrifugation at 13000 rpm for 10 min at 4°C by a table-top centrifuge and further purified by chromatography on a Mono Q HR 5/5 column (GE Healthcare) with a linear gradient of 0.1-1.0 M sodium chloride in chromatography buffer (20 mM HEPES–KOH pH 7.6, 10 μM zinc chloride, 1 mM EDTA pH 8.0, and 20% glycerol). Proteins were dialyzed against dialysis buffer (20 mM HEPES–KOH pH 7.6, 500 mM potassium acetate, 10 μM zinc chloride, 1 mM EDTA pH 8.0, 20% glycerol, 1 mM DTT, and 1× protease inhibitors). 1-5 μM SNAP_f_ fused activator was incubated in a 1× PBS solution containing up to threefold excess of SNAP-surface 649 (New England BioLabs, #S9159S), 1 mM DTT, and 1× protease inhibitors at 4°C for 1.5 hr on a rotator in the dark. Labeled activators were incubated with Ni^2+^-NTA-agarose resin (Goldbio) at 4°C for 1 hr on a rotator. The protein-bound resin was extensively washed with 1× PBS supplemented with 10% glycerol, 10 μM zinc chloride, 1 mM DTT, and 1× protease inhibitors. Labeled proteins were eluted with 300 mM imidazole in 1× PBS supplemented with 10% glycerol, 10 μM zinc chloride, 1 mM DTT, and 1× protease inhibitors and stored at -80°C. Protein purify was monitored by SDS-PAGE and Coomassie blue staining. Labeling of SNAP_f_ fused activators and residual dye depletion were confirmed by in-gel fluorescence imaging on a Typhoon imager (GE Healthcare).

### Preparation of naked and chromatinized template DNAs

Biotinylated, dye-labeled DNA template (**Figure 1A** and **Table S1**) was prepared by PCR from pUC18-G5CYC1 G- (**Table S1**) with Platinum Taq DNA polymerase (Invitrogen) and primers Biotin-universal and AF488-M13 rev2 (**Table S1**). The PCR product was purified using DNA SizeSelector-I SPRI magnetic beads (Aline Biosciences) according to the manufacturer’s instruction.

To make chromatin with unmodified nucleosomes, recombinant yeast histone octamers were purified from bacteria as previously described (Migl et al., 2020). Plasmid pDMM140 pScH3 Oct (**Table S1**) was transformed into Rosetta 2 *E. coli* cells (Novagen). Transformed cells were grown at 37°C in LB or 2X YT + glucose media with appropriate antibiotics to an optical density at 600 nm of 0.6-1.0. Cells were induced with 0.25-0.5 mM IPTG and grown overnight at 18°C. Cells were pelleted, resuspended in 120 ml of high salt buffer (HSB: 50 mM HEPES pH 7.5, 2 M sodium chloride, 10% glycerol, 1 mM TCEP) containing protease inhibitors (aprotonin, pepstatin, leupeptin, and PSMF), and lysed by sonication. The lysate was centrifuged at 40,000 x *g* for 1 hr at 4°C, and the supernatant was incubated with 4 ml TALON metal affinity resin (Clontech) at 4°C for 1 hr. The protein-bound resin was extensively washed with HSB and, the protein was eluted with HSB containing 50 mM EDTA pH 8.0 and 400 mM imidazole. The eluate was concentrated in an Amicon centrifugal filter (molecular weight cut-off, 10 kDa) by repeated centrifugations (3000 *x g* for 15 min at 4°C). The concentrated histone octamers were further purified on a 120 ml Superdex 200 size exclusion column (GE Healthcare) with SEC buffer (30 mM HEPES-KOH pH 7.5, 2 M sodium chloride, 10% glycerol, 1 mM TCEP).

Semi-synthetic human histones (unmodified or with fully defined native tetra-acetylations at H3 [K4,K9,K14,K18] or H4 [K5,K12,K16,K20]) were synthesized and assembled into octamers as previously described (Grzybowski et al., 2015). Briefly, recombinant histones lacking N-terminal tails were processed to generate an N-terminal cysteine, which was then used for ligation to tail peptides carrying the specified modifications. Desulfurization produces a scarless junction. Octamers were reconstituted by co-renaturation of all four histones, followed by size exclusion chromatography.

For nucleosome assembly, yeast histone chaperone Nap1 and Isw1a complex were purified as previously described (Joo et al., 2019a; Joo et al., 2019b). For the purification of yeast Nap1, pGEX-6P-1-NAP1 plasmid (**Table S1**) was transformed into BL21 (Codon+, DE3) cells. Cells were grown at 37°C in LB medium with appropriate antibiotics to an optical density at 600 nm of 0.5. Proteins were induced with 0.2 mM IPTG for 4 hr at 30°C. Cells were harvested by centrifugation in a Sorvall GS3 or SLA-3000 rotor at 5000 rpm for 10 min at 4°C, resuspended in 1× PBS, and lysed by sonication. The soluble extract was isolated after centrifugation in a GSA rotor at 13000 rpm for 1 hr at 4°C and incubated with Glutathione agarose resin (Pierce 25237) for 1 hr on a rotator at 4°C. The protein-bound resin was extensively washed with 1× PBS, and bound proteins were eluted by digestion with 20 units of Prescission protease (GE Healthcare) in 0.5 ml of Prescission protease cleavage buffer (50mM Tris-HCl pH 7.0, 150 mM sodium chloride, 1 mM EDTA pH 8.0, 1 mM DTT) at 4 °C for 16 hr. The eluate was further purified by chromatography on a Mono Q HR 5/5 column (GE Healthcare) with a linear gradient of 0.1-1.0 M sodium chloride in Nap1 chromatography buffer (20 mM Tris-Cl pH 7.5, 0.5 mM EDTA pH 8.0, 10% glycerol, 1 mM DTT, and 0.1 mM PMSF). Nap1 fractions were combined and dialyzed against Buffer R (10 mM HEPES–KOH pH 7.6, 10 mM potassium chloride, 1.5 mM magnesium chloride, 0.5 mM EGTA pH 8.0, 10% glycerol, 2.5 mM β-glycerophosphate, 1 mM DTT, and 0.2 mM PMSF).

For the purification of yeast Isw1a complex, strain YF2164 (**Table S1**) was grown at 30°C in 6 liters of YPD to an optical density at 600 nm of 2.3. Cells were washed with distilled water, pelleted by centrifugation in a Sorvall GS3 or SLA-3000 rotor at 5000 rpm for 10 min at 4°C, and resuspended in TAP lysis buffer (50 mM Tris pH 8.0, 300 mM potassium chloride, 1 mM PMSF, 1 μg/ml benzamidine and 1× protease inhibitors). Cells were lysed by bead beating with pre-chilled glass-beads (0.5 mm). The lysate was centrifuged in a Sorvall GS3 or SLA-3000 rotor at 12000 rpm for 15 min at 4°C, and the supernatant was incubated with 0.7 ml of IgG Sepharose beads (GE Healthcare) at 4°C for 3.5 hr. The protein bound resin was extensively washed with TAP lysis buffer and then washed with TEV cleavage buffer (10 mM Tris-Cl pH 8.0, 300 mM potassium chloride, 1 mM magnesium chloride, 10% glycerol, 0.1% NP-40, 1 mM DTT, 1 μg/ml benzamidine, and 1× protease inhibitors). Proteins were eluted by digestion with 600 units of recombinant TEV protease (Lucast et al., 2001) in 3 ml of TEV cleavage buffer overnight at 4 °C. The eluate was incubated with 0.3 ml of Calmodulin Sepharose 4B beads (GE Healthcare) in the addition of 7 ml of Calmodulin binding buffer (10 mM Tris-Cl pH 8.0, 300 mM potassium chloride, 1 mM magnesium acetate, 2 mM calcium chloride, 1 mM imidazole, 10% glycerol, 0.1% NP-40, 10 mM β-mercaptoethanol, 1 mM PMSF, 1 μg/ml benzamidine, and 1× protease inhibitors) at final concentration of 10 mM CaCl_2_ for 3 hr at 4 °C. The protein bound Calmodulin resin was extensively washed with Calmodulin binding buffer, and Isw1a complex was eluted with Calmodulin elution buffer (10 mM Tris-Cl pH 8.0, 300 mM potassium chloride, 1 mM magnesium acetate, 1 mM imidazole, 3 mM EGTA, 10% glycerol, 0.1% NP-40, 1 mM PMSF, 1 μg/ml benzamidine, and 1× protease inhibitors).

Chromatinized templates were assembled by enzymatic reconstitution as previously described (Joo et al., 2019b). Yeast histone octamer, 2 μg of Nap1, and 30 ng of Isw1a complex were mixed in 90 ul of Buffer R adjusted to 50 mM potassium chloride and incubated for 30 min on ice. The reaction was then brought to 110 μl total reaction volume by addition of 660 ng of the naked DNA template together with final concentrations of 3 mM ATP, 4.2 mM magnesium chloride, 700 ng of creatine phosphokinase, and 30 mM phosphocreatine. The assembly reaction was incubated for 5 hr at 30°C and stored at 4°C until use within 24 hr. Optimal DNA:histone ratio and nucleosome assembly of chromatinized template were confirmed by a partial MNase digestion assay as previously described (**Figure 4B**) (Fyodorov and Kadonaga, 2003).

### *In vitro* transcription

*In vitro* transcription assays were conducted as previously described (Joo et al., 2019b; Sikorski et al., 2012). 250 ng of plasmid pUC18-G5CYC1 G-(SB649) (**Table S1**), 325 nM activator, 10 mM phosphocreatine, 0.1-0.2 units of creatine kinase, 0.33 units of RNasin (Promega), and 10 μl of nuclear extract (25 - 40 μg of protein) were incubated in ATB buffer (20 mM HEPES-KOH pH 7.6, 100 mM potassium acetate, 1 mM EDTA pH 8.0, 5 mM magnesium acetate) for 5 min at room temperature. 500 μM ATP, 500 μM CTP, and 20 μM UTP along with 0.3 μCi of α-^32^P UTP (PerkinElmer) were added to the mixture to initiate transcription. After 45 min incubation at room temperature, transcripts were recovered by phenol-chloroform extraction and ethanol precipitation, separated by gel electrophoresis (8M urea-6% polyacrylamide gel), and analyzed by phosphoimaging on a Typhoon imager (GE healthcare).

### Single-molecule microscopy of eukaryotic transcription reactions

Single-molecule fluorescence microscopy experiments were performed as previously described with minor modifications (Baek et al., 2021). A multi-wavelength micromirror TIRF microscope system (Friedman et al., 2006) was operated using the custom software Glimpse, implemented with LabView (National Instruments; Austin, TX) (https://github.com/gelles-brandeis/Glimpse). Passivated slides, flow chambers, and fiducial bead markers (Molecular Probes T-10711) were prepared as described in (Rosen et al., 2020). 10 pM of the biotinylated/AF488-labeled templates (e.g. naked and chromatinized) were immobilized sequentially on the same slide surface through biotin-streptavidin in TPA buffer (50 mM Tris-acetate pH 7.9, 100 mM potassium acetate, 8 mM magnesium acetate, 27 mM ammonium acetate, and 1 mg/ml bovine serum albumin (BSA)) supplemented with the oxygen scavenging system (0.9 units/ml protocatechuate dioxygenase (Sigma P8297) and 5 mM protocatechuic acid (Sigma 03930590)) and triplet state quenchers (1 mM propyl gallate (Sigma 02370), 2 mM 4-nitrobenzyl alcohol (Sigma N12821), and 1 mM Trolox (Sigma 238813)). Two template images, one with the fluorescent spots of the first template and the other with fluorescent spots of both templates, were acquired using the 488 nm excitation laser at 1.2 mW laser power. After the template immobilization, transcription reactions containing 5-10 mg protein/ml yeast nuclear extract, indicated concentrations of Gal4-SNAP_f_ or Gal4-SNAP_f_-VP16, 400 μM 4 NTPs (unless otherwise noted), 2 mM acetyl-CoA (unless otherwise noted), 20 mM HEPES-KOH pH7.6, 100 mM potassium acetate, 20 μg/ml *E. Coli* genomic DNA, 5 mM magnesium acetate, 1 mM EDTA pH 8.0, oxygen scavenging reagents, and triplet state quenchers were introduced in a flow chamber at room temperature. In the experiments carried out in the absence of 4 NTPs, an ATP depletion system, 20 mM glucose and 2 units hexokinase (Sigma H4502), was added instead of 4 NTPs. Images of DY649-labeled activator and DY549-labeled SAGA were collected individually by alternating the 532 nm and 633 nm excitation lasers (0.5 s/frame for each excitation laser) at 1.2 mW and 600 μW laser powers, respectively. The 785 nm excitation laser was used to maintain focus automatically.

### General image analysis procedure

Image data analysis was performed with custom software imscroll, implemented with MATLAB (The MathWorks; Natick, MA), as previously described (Baek et al., 2021; Friedman and Gelles, 2015; Friedman et al., 2013)(https://github.com/gelles-brandeis/CoSMoS_Analysis). To correct for chromatic aberration and other sources of spatial displacement between the three laser channels, a separate mapping experiment was done with surface-tethered oligonucleotides containing three fluorescent dyes (AF488, Cy3, and Cy5) in each molecule. Hundreds of calibration spot pairs between different color channels (e.g., blue versus green) were imaged and mapped, and the resulting matrix was applied to the images collected during the transcription reaction (Friedman and Gelles, 2015).

Transcription reactions typically contained two DNA templates, requiring sequential imaging (see above). Fluorescent spots were automatically selected by imscroll. Those observed both before and after immobilizing the second template mapped the locations of the first immobilized template. Fluorescent spots observed only on the image acquired after second template immobilization were mapped as second template locations. Closely spaced template pairs were removed to avoid confusing signals. Automatically selected areas with no fluorescent spots on both images were mapped as off-target locations.

To correct for spatial stage drift during the experiment, each field of view contained several immobilized fluorescent beads as fiducial markers (Invitrogen, T10711). Their trajectories were mapped in each channel and the corrections applied to the protein images. For protein mapping, fluorescence intensity within small regions (0.16 or 0.28 μm^2^; 3×3 or 4×4 pixels, respectively) that overlapped the pre-defined template locations were integrated to produce fluorescence intensity time records at each template location (**Figures 2A, 5B, S1, and S1A**). Fluorescence intensities of proteins at each template location were converted to define a bound (presence of a fluorescent spot) or unbound state (absence of a fluorescent spot) by thresholding two characteristics, distance from the center of template location and spot intensity. These binary-coded states were visualized as rastergrams using custom MATLAB scripts (**Figures 2C, 3A, 5A, 6A and B, 7C, S1B, S1A and B, and S1F**).

## QUANTIFICATION AND STATISTICAL ANALYSIS

### Time to first binding plots and association rate constants

The unbound “absent” intervals preceding the initial observed activator or SAGA binding to each template were sorted by length and plotted as cumulative time-to-first binding curves (**Figures 2B, 4C, 7A and E, S1A, S1B, S1A, S1A and C**). Template-specific apparent first-order rate constants of activator and SAGA (*k*_on_ values in **Figures 2D, 3B, 6C, 7B and F**) were determined by curve fitting as previously described (Friedman and Gelles, 2015). Association rates calculated from these initial unbound intervals avoid complications from photobleaching and/or photoblinking effects that might affect accuracy in counting subsequent binding events. The CoSMoS data analysis accounts for both nonspecific factor binding at off-target locations, as well as a fraction of unused templates. Nonspecific association rate constants (*k*_ns_) are calculated using off-target locations. The active fraction is defined as follows: *A*_f_ ≤ 1; a faction of template locations is capable of factor binding. The template-specific apparent first-order rate constants (*k*_on_) are derived from a model that assumes two modes of factor binding, (1) factor binding at the active fraction of target locations at a rate that includes both specific and non-specific binding (*k*_on +_ *k*_ns_) and (2) factor binding on an inactive fraction (1 - *A*_f_) at the rate *k*_ns_. Standard errors of rate constants are determined by bootstrapping from 2000 iterations. Fit parameters are listed in **Table S1**.

### Template-specific binding frequency of Spt7

At lower activator concentrations, a large fraction of SAGA binding events occurred before activator arrival. To better quantitate the effect of activator on Spt7 recruitment, we used a second method to calculate the template-specific frequency of Spt7 arrival when activator was either present or absent. First, the total time course for each template or off-target location was sorted into intervals when fluorescent activator was present or absent. Total binding frequency of Spt7 within each class was calculated by dividing the number of Spt7 binding events by the sum of time intervals during which the SAGA complex was absent. The template-specific binding frequency of Spt7 was determined by subtracting the corresponding frequency of Spt7 binding to off-template locations from the total frequency of Spt7 binding to on-template locations. (**Figures 2E, 4D, S1D, and S1E**). Standard errors were determined from the standard deviation of a binomial distribution, the uncertainty in frequency 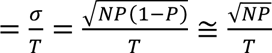 (*if P is close to* 0 *or* 1), where *N* is the number of image frames, *P* is the probability of a binding event per frame, and *T* is the total time during which the target site is empty.

### Cumulative distributions and surviving fractions of factor dwell times

To compare the duration of activator or SAGA binding in different conditions, cumulative template-specific binding frequency distributions (**Figure S1B**) or template-specific surviving fractions (**Figures 2F, 3C and D, 4E, 6D, 7D, S1, S1B and D**) were plotted. Note that “bound” time intervals mark when one or more molecules of a factor are present on the template, so if binding of individual molecules overlap, the dwell times measured can overestimate single molecule binding duration. Conversely, binding frequencies do not include events where a molecule binds a template already bound by another similar molecule, thereby underestimating binding frequency. Cumulative binding frequency distributions of factors are obtained by dividing the number of occupancy events with dwell times longer than the x-axis time point by the sum of times when the factor is absent. Similarly, surviving factions of factors were determined by taking the number of binding events with longer dwell time than the x-axis time point and dividing by the total number of binding events. For template-specific cumulative binding frequency distributions and surviving fractions, cumulative frequency of off-template locations or the number of off-template binding events at each dwell time point were subtracted from those of on-template locations, respectively. The 90% confidence interval was determined by bootstrapping. A maximum likelihood algorithm using only the list of dwell intervals was used to calculate fit parameters in Figure S1B as previously described (Friedman and Gelles, 2015).

### Analyzing orders of factor arrival during overlapped binding of activator and SAGA

To detect correlations between activator and SAGA arrivals, intervals having overlapping binding of both factors on the same template were extracted. To avoid any potential artifacts from photoblinking, if multiple bindings were seen for one of the two factors, only the initial binding time was used. Probability density histograms of the delay times between arrivals of activator and SAGA (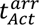−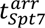) were plotted (**Figure 5C**). These times can be divided into three categories; activator-first, SAGA-first, and simultaneous within the experimental time resolution (± 1.4 seconds approximately). For activator-first and the SAGA-first categories, histogram bin widths were adjusted so that each bin has roughly the same number of counts. Standard errors of bar heights were determined by the binomial distribution. To account for the possible random coincidence of activator and SAGA binding on the same template, intervals of activator from each template location were randomly paired with SAGA intervals at a different template location. The scrambled data generated in this way was then subject to the same analysis.

## Author contributions

J. Jeon did the experiments and data analysis, with assistance from L. Friedman and D. Seo. Conception and experimental design was done by S. Buratowski, J. Gelles, and J. Jeon. A. Adeleke, B. Graham, and E. Patteson created the acetylated and unmodified human nucleosomes. The paper was written by J. Jeon and S. Buratowski, with additional editing by L. Friedman and J. Gelles.

## Acknowledgments

We are grateful to Inwha Baek for advice on CoSMoS; Sarah Le for creating the Gal4-SNAP-Rap1 activator; David Migl for histone expression vectors, Song Tan (Penn State) and Phil Cole for advice on generating modified nucleosomes, and to Dan Zhou for advice and Matlab scripts for CoSMoS analysis. Michael Keogh (EpiCypher) was invaluable for providing modified nucleosomes and comments on the manuscript. This work was supported by NIH R01 CA246500 to S.B. and J.G.

## SUPPLEMENTARY FIGURE LEGENDS

**Figure S1.**
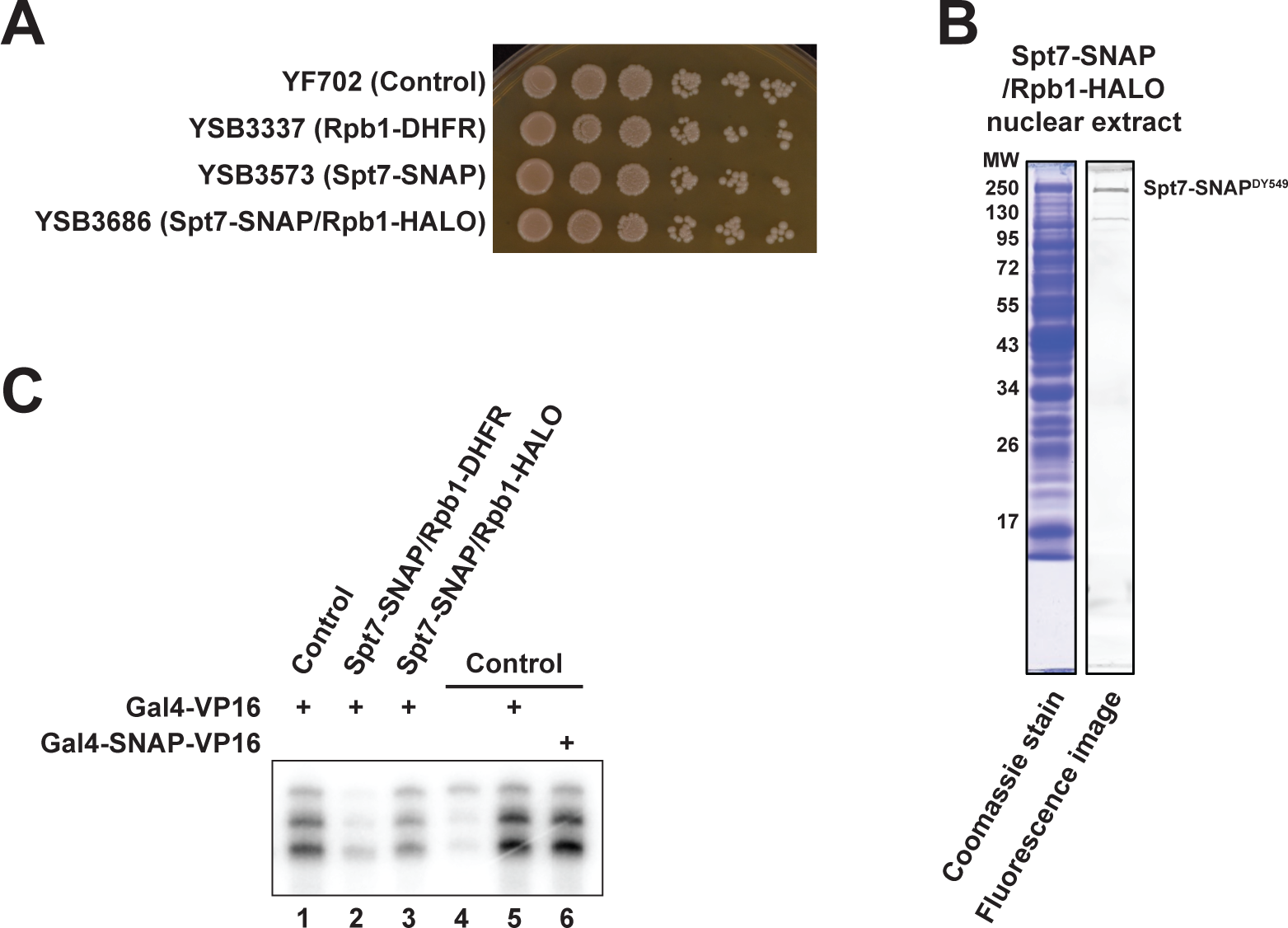
Validation of Spt7-tagged yeast strains. **A.** Spotting assay for cell growth. Four-fold serial dilutions of the untagged parent strain (YF702) and strains carrying the indicated fusions (YSB3337, YSB3573, and YSB3686) were grown on a YPD plate at 30°C. No significant differences in cell growth were seen. **B.** Coomassie staining and fluorescence imaging of an SDS-PAGE gel resolving nuclear extract from strain YSB3686 (Spt7-SNAP ^DY549^/Rpb1-HALO). **C.** Bulk *in vitro* transcription assay with yeast nuclear extracts, transcription activator Gal4-VP16, and plasmid template. Radioactive 32P transcription products were separated on an 8 M urea-8% polyacrylamide gel and analyzed by phosphoimager. Nuclear extracts from labeled fusion strains were slightly less active than untagged YF702 cells (lane 1-3), at least in part due to the extra time and manipulations needed for fusion protein labeling and removal of free dye. Transcription activation by Gal-VP16 was not affected by insertion of the SNAP_f_ tag between the Gal4 DNA binding domain and VP16 activation domain (compare lanes 5 and 6 to lane 4).

**Figure S2.**
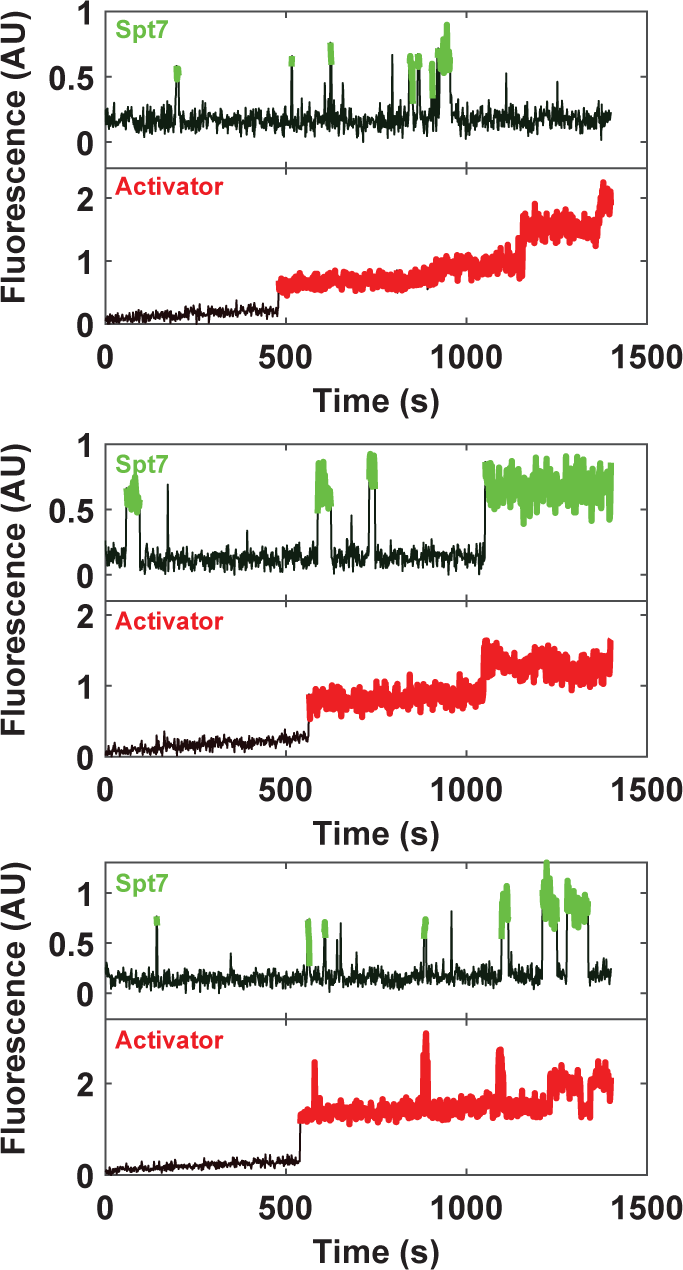
Time records of SAGA and activator fluorescence intensity on three different naked templates in the presence of 10 nM Gal4-SNAP^DY649^-VP16. Spt7-SNAP^DY549^ (green) or activator (Gal4-SNAP^DY649^-VP16, red). Periods of fluorescent protein colocalization with the DNA template are colored. The records illustrate more frequent and longer duration SAGA binding events when activator is present on the same template.

**Figure S3.**
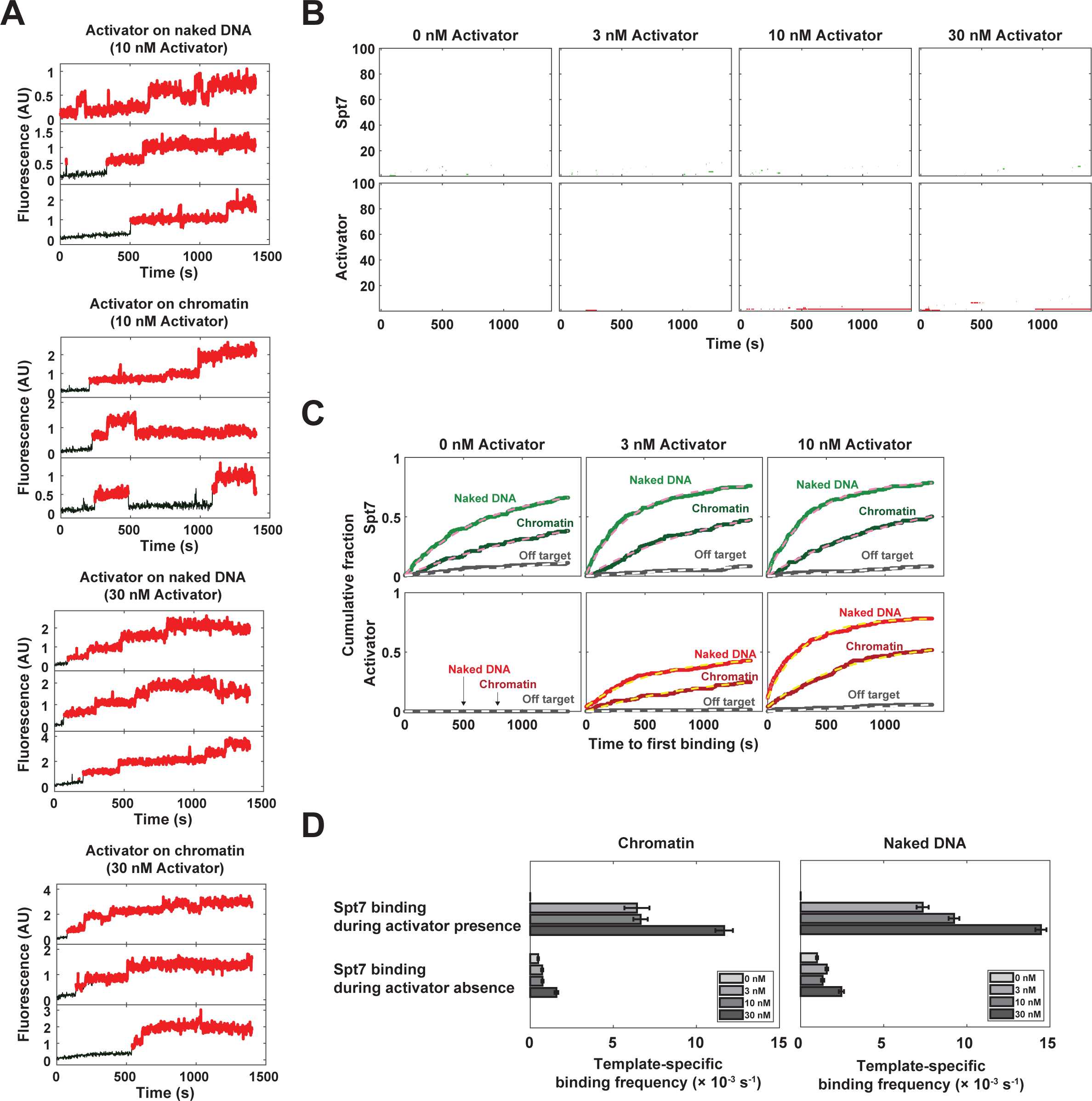
A. Representative time records of Gal4-SNAP^DY649^-VP16 activator on naked and chromatinized templates at 10 nM and 30 nM activator concentrations. Note the intensity jumps reflecting multiple activator molecules binding to the five Gal4 binding site UAS. **B.** Rastergrams of Spt7-SNAP^DY549^ (green) and Gal4-SNAP^DY649^-VP16 (activator, red) binding to “off-target” areas of the slide where no template DNA is bound. 100 randomly chosen off-target locations show only sparse activator and Spt7 binding events at various activator concentrations (0, 3, 10 and 30 nM). **C**. Cumulative time-to-first-binding distributions for Spt7-SNAP^DY549^ (upper panels) and Gal4-SNAP^DY649^-VP16 (lower panels) on naked (light colors) versus chromatin (dark colors) templates at various activator concentrations (0, 3, and 10 nM). Gray curves show off-target background binding to slide surface, and dashed lines indicate curve fits. The corresponding data for 30 nM activator appears in Figure 4C. **D.** Total SAGA binding frequencies on chromatinized (left panel) and naked (right panel) templates when activator is present or absent. Four different activator concentrations (0, 3, 10 and 30 nM as indicated in key) were tested. Error bars showing standard error were determined by the binomial distribution. Note that at 0 nM activator there is no “activator presence”, so the total frequency of Spt7 binding is “during activator absence”.

**Figure S4.**
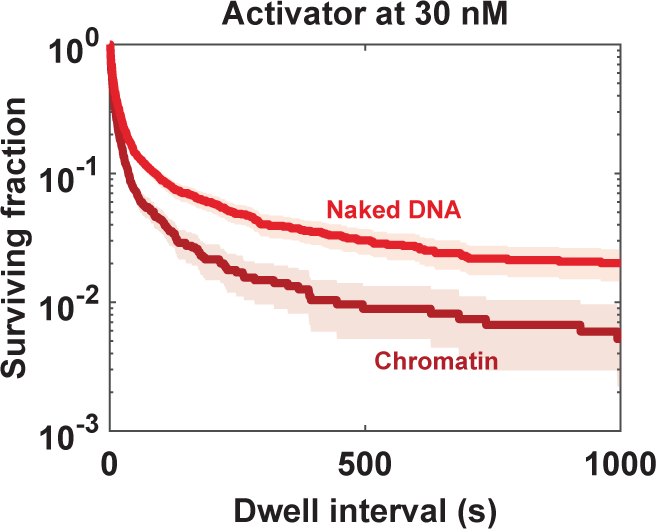
Survival plot of Gal4-SNAP^DY649^-VP16 dwell intervals on naked (light red) and chromatinized (dark red) templates at 30 nM concentration. Shaded areas indicate the 90% confidence interval as determined by bootstrapping.

**Figure S5.**
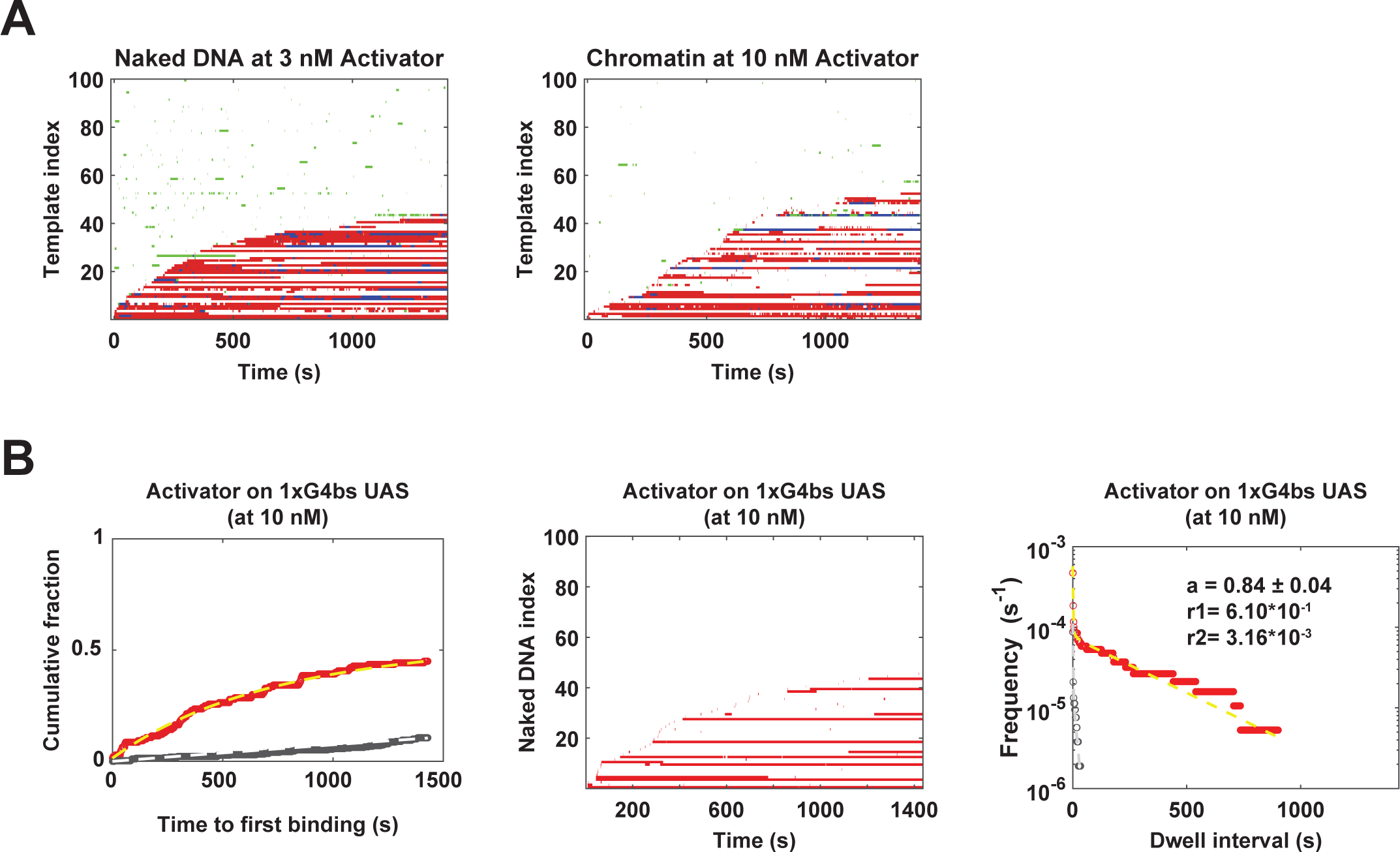
Long duration activator binding correlates with long duration SAGA binding. **A.** Dual-color rastergrams of Gal4-SNAP^DY649^-VP16 and Spt7-SNAP^DY549^ binding events. Activator binding records from 100 randomly chosen naked (left panel) and chromatinized (right panel) template locations were binary-coded (red color indicates colocalization with DNA) and sorted by time to first activator binding, from bottom to top. Next, corresponding binary-coded SAGA binding records (green) from the same template were overlaid. Simultaneous binding of SAGA and activator was colored as blue. Note that different activator concentrations (3 nM Gal4-SNAP^DY649^-VP16 for naked and 10 nM Gal4-SNAP^DY649^-VP16 for chromatinized) were used for dual rastergram analyses to better match the activator binding occupancies of the two template types. **B.** Gal4-SNAP^DY649^-VP16 (10 nM) binding to naked DNA templates containing a single Gal4 binding site (1xG4bs), in the absence of NTPs. Left panel shows cumulative time-to-first-binding distribution of Gal4-SNAP^DY649^-VP16 on the templates (red) or off-targets (gray). Dashed lines are single exponential curve fits. Middle panel displays rastergram of Gal4-SNAP^DY649^-VP16 activator binding (red) records from 100 randomly chosen templates. Right shows the cumulative distribution plots of Gal4-SNAP^DY649^-VP16 dwell intervals on the single Gal4 UAS containing template (red) or at off-target locations (gray dotted line). Dashed lines show biexponential distribution curve fitting for the DNA locations (yellow; *a* = 0.84 ± 0.04 *r_1_* = (6.1 ± 0.6) × 10^-1^ s^-1^, *r_2_* = (3.16 ± 1.34) × 10^-3^ s^-1^) and the off-target locations (light gray; *a* = 0.86 ± 0.06, *r_1_* = (7.7 ± 0.6) × 10^-1^ s^-1^, *r_2_* = (6.9 ± 2.4) × 10^-2^ s^-1^).

**Figure S6.**
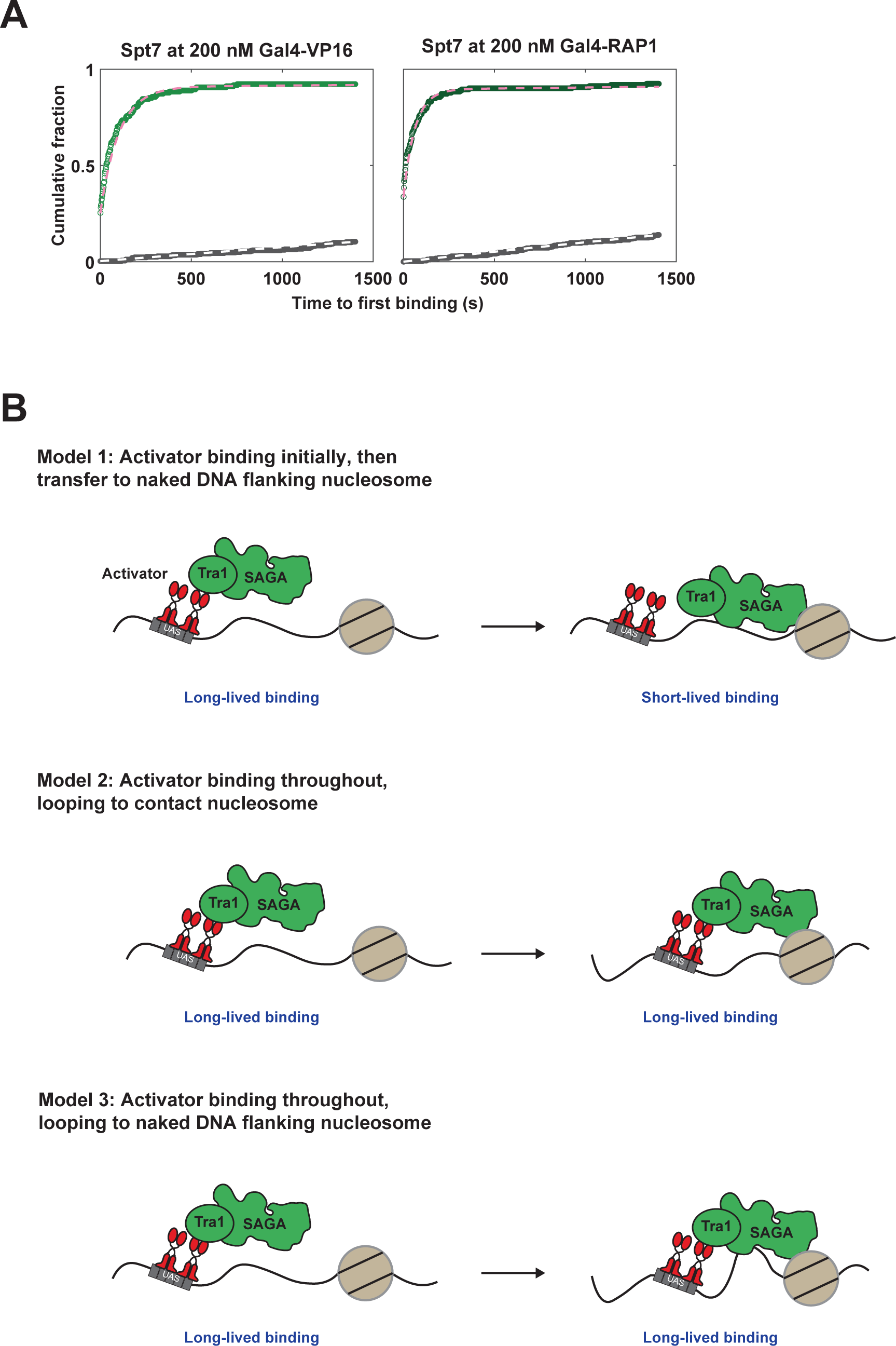
A. Cumulative time-to-first-binding distributions for Spt7-SNAP^DY549^ on naked DNA templates, in the absence of NTPs. Gal4 DNA binding domain derivatives fused to either the VP16 (light green, left) or Rap1 (dark green, right) activation domains were used. Gray curves illustrate off-target background binding to slide surface, and dashed lines indicate curve fits. Note that these curves correspond to the full data set represented by the 100 randomly chosen DNAs used for the rastergrams in Figure 6B. B. Three models for how activators can “recruit” SAGA HAT activity to promoters. In all cases, SAGA is initially tethered to the UAS/enhancer via interactions with the activation domain. In Model 1, SAGA transfers to nearby naked DNA and is released from the activator. Acetylation occurs at nucleosomes flanking the naked DNA. In Model 2, SAGA remains tethered to the activator while the HAT activity targets nearby nucleosomes without any additional DNA contact. In Model 3, SAGA remains tethered to the activator, but also contacts nearby naked to target the flanking nucleosomes.

**Figure S7.**
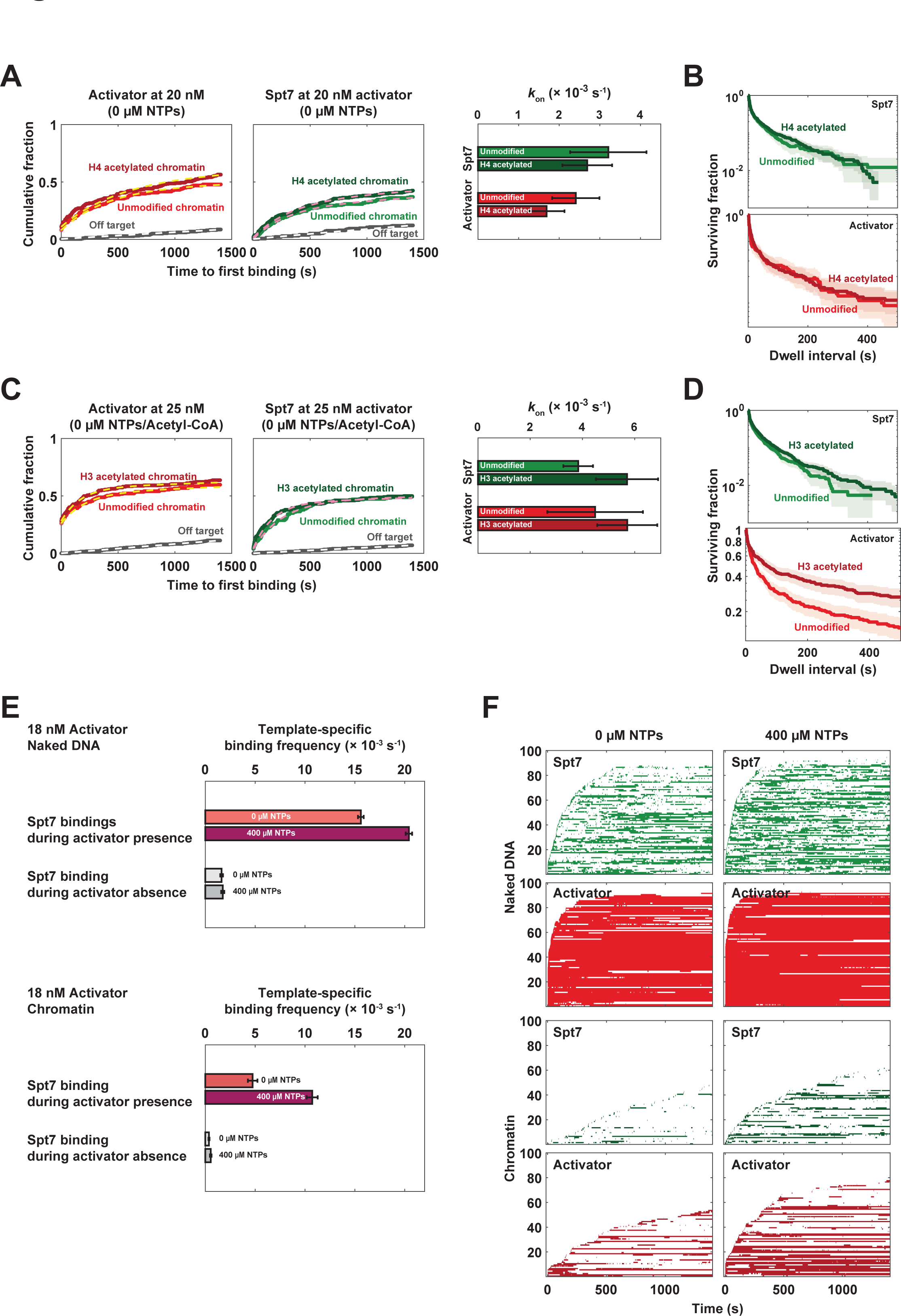
Histone acetylation and NTPs affect both SAGA and activator binding to chromatin templates. **A.** Spt7-SNAP^DY549^ (green) and Gal4-SNAP^DY649^-VP16 (red) bindings were simultaneously imaged on transcription templates chromatinized with human histone octamers containing either unmodified (light colors) or tetra-acetylated (dark colors) histone H4. Left and middle panels show cumulative time-to-first-binding curves. Right panel shows the apparent binding rates of Spt7-SNAP^DY549^ and Gal4-SNAP^DY649^-VP16 activator derived from curve fitting to a single exponential function. Compare these results in the absence of NTPs to those in the presence of NTPs shown **in** Figures 7A and B. **B.** Survival plots for Spt7-SNAP^DY549^ and Gal4-SNAP^DY649^-VP16 dwell intervals on unmodified versus H4 acetylated chromatin templates in the absence of NTPs. Compare to Figure 7D results obtained in the presence of NTPs. Shaded regions indicate 90% confidence intervals determined by bootstrapping. **C** and **D.** Same as panels **A** and **B,** except for utilizing H3 tetra-acetylated chromatin instead of H4 tetra-acetylated chromatin. This experiment was conducted in the absence of NTPs and acetyl-CoA. **E.** SAGA binding frequencies on naked (top panel) and chromatinized (bottom panel) templates when activator is present or absent. Both -NTPs and +NTPs conditions were tested. Error bars were determined by the binomial distribution. **F.** Rastergrams of Gal4-SNAP^DY649^-VP16 (red) and Spt7-SNAP^DY549^ (green) binding on naked DNA, chromatinized template, and off-target locations at 18 nM activator in the presence or absence of NTPs.

## Supplementary tables

**Table S1:** *S. cerevisiae* strains used in this study.

| Strain | Genotype | Source | Target proteins |
| --- | --- | --- | --- |
| YF702/CB012 | MATa, ura3-1, leu2-3,112, trp1-1, his3-11,15, ade2-1, pep4Δ::HIS3, prbΔ::his3, prc1Δ::hisG | (Inada et al., 2002) | Wild-type |
| YSB3337 | MATa, ura3-1, leu2-3,112, trp1-1, his3-11,15, ade2-1, pep4Δ::HIS3, prbΔ::his3, prc1Δ::hisG, RPB1-DHFR::Hygromycin<sup>R</sup> | (Rosen et al., 2020) | Rpb1<sup>DHFR</sup> |
| YSB3562 | MATa, ura3-1, leu2-3,112, trp1-1, his3-11,15, ade2-1, pep4Δ::HIS3, prbΔ::his3, prc1Δ::hisG, RPB1-DHFR::Hygromycin<sup>R</sup>, SPT7-HA<sub>3</sub>-SNAP<sub>F</sub>:: KANMX6 | This study | Rpb1<sup>DHFR</sup><br>Spt7<sup>SNAP<sub>F</sub></sup> |
| YSB3573 | MATa, ura3-1, leu2-3,112, trp1-1, his3-11,15, ade2-1, pep4Δ::HIS3, prbΔ::his3, prc1Δ::hisG, SPT7-HA<sub>3</sub>-SNAP<sub>F</sub>:: KANMX6 | This study | Spt7<sup>SNAP<sub>F</sub></sup> |
| YSB3685 | MATa, ura3-1, leu2-3,112, trp1-1, his3-11,15, ade2-1, pep4Δ::HIS3, prbΔ::his3, prc1Δ::hisG, SPT15-HA<sub>3</sub>-SNAP<sub>F</sub>:: Hygromycin<sup>R</sup>, SPT7-HA<sub>3</sub>-HALO::NATMX | This study | Spt15<sup>SNAP<sub>F</sub></sup><br>Spt7<sup>HALO</sup> |
| YSB3686 | MATa, ura3-1, leu2-3,112, trp1-1, his3-11,15, ade2-1, pep4Δ::HIS3, prbΔ::his3, prc1Δ::hisG, RPB1-HA<sub>3</sub>-HALO::NATMX, SPT7-HA<sub>3</sub>-SNAP<sub>F</sub>::KANMX6 | This study | Rpb1<sup>HALO</sup><br>Spt7<sup>SNAP<sub>F</sub></sup> |
| YF2164 | MATa, ura3-52, leu2-3,112, trp1-1, his3-11,15, ade2-1, can1-100, pep4Δ::LEU2, IOC3-TAP::TRP1 | (Johnson et al., 2009) | Purification of loc3-TAP/Isw1 complex |

**Table S2:**
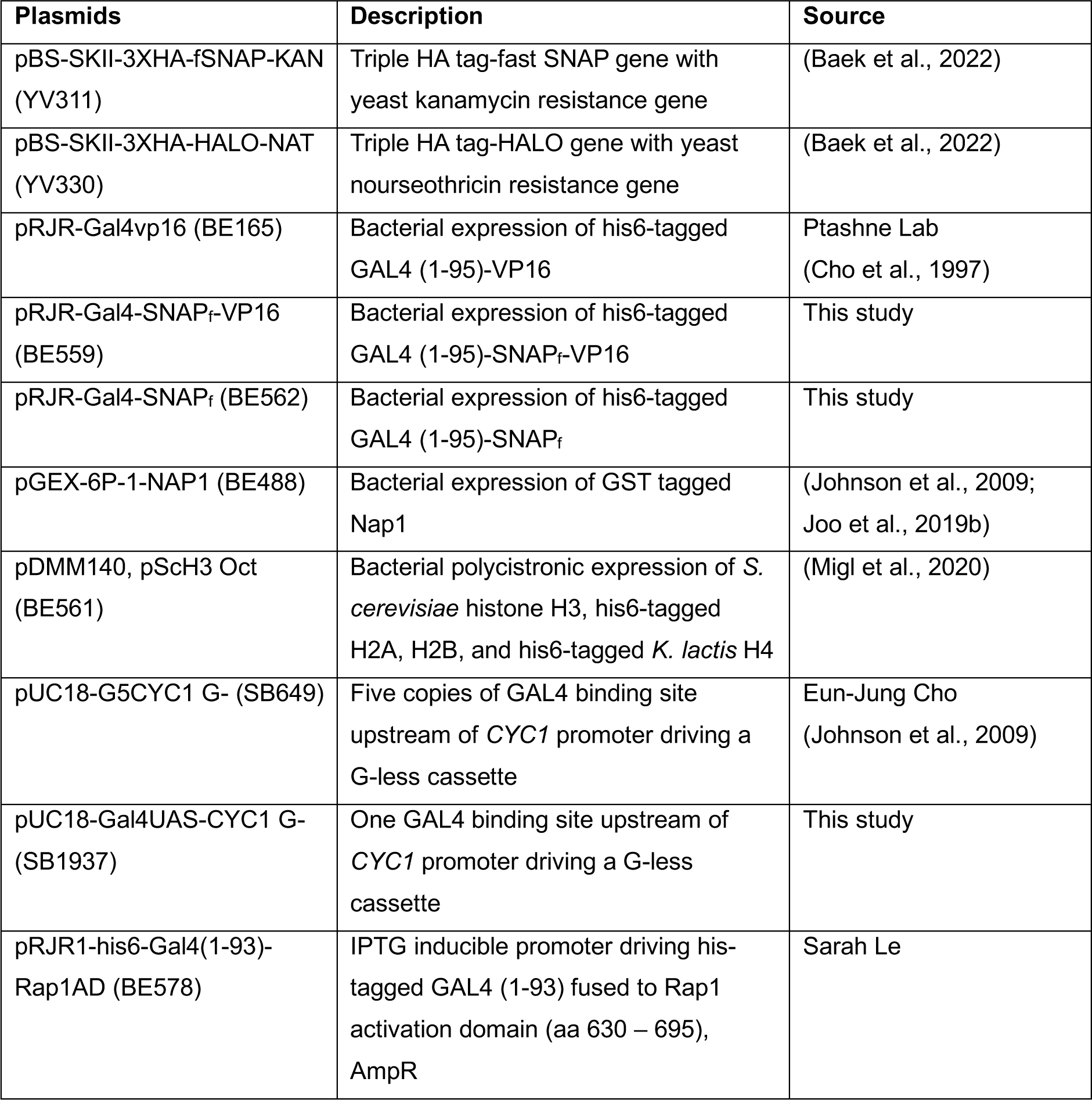
Plasmids used in this study.

| Plasmids | Description | Source |
| --- | --- | --- |
| pBS-SKII-3XHA-fSNAP-KAN (YV311) | Triple HA tag-fast SNAP gene with yeast kanamycin resistance gene | (Baek et al., 2022) |
| pBS-SKII-3XHA-HALO-NAT (YV330) | Triple HA tag-HALO gene with yeast nourseothricin resistance gene | (Baek et al., 2022) |
| pRJR-Gal4vp16 (BE165) | Bacterial expression of his6-tagged GAL4 (1-95)-VP16 | Ptashne Lab (Cho et al., 1997) |
| pRJR-Gal4-SNAP<sub>f</sub>-VP16 (BE559) | Bacterial expression of his6-tagged GAL4 (1-95)-SNAP<sub>f</sub>-VP16 | This study |
| pRJR-Gal4-SNAP<sub>f</sub> (BE562) | Bacterial expression of his6-tagged GAL4 (1-95)-SNAP<sub>f</sub> | This study |
| pGEX-6P-1-NAP1 (BE488) | Bacterial expression of GST tagged Nap1 | (Johnson et al., 2009; Joo et al., 2019b) |
| pDMM140, pSch3 Oct (BE561) | Bacterial polycistronic expression of *S. cerevisiae* histone H3, his6-tagged H2A, H2B, and his6-tagged *K. lactis* H4 | (Migl et al., 2020) |
| pUC18-G5CYC1 G- (SB649) | Five copies of GAL4 binding site upstream of *CYC1* promoter driving a G-less cassette | Eun-Jung Cho (Johnson et al., 2009) |
| pUC18-Gal4UAS-CYC1 G- (SB1937) | One GAL4 binding site upstream of *CYC1* promoter driving a G-less cassette | This study |
| pRJR1-his6-Gal4(1-93)-Rap1AD (BE578) | IPTG inducible promoter driving his6-tagged GAL4 (1-93) fused to Rap1 activation domain (aa 630 – 695), AmpR | Sarah Le |

**Table S3:**
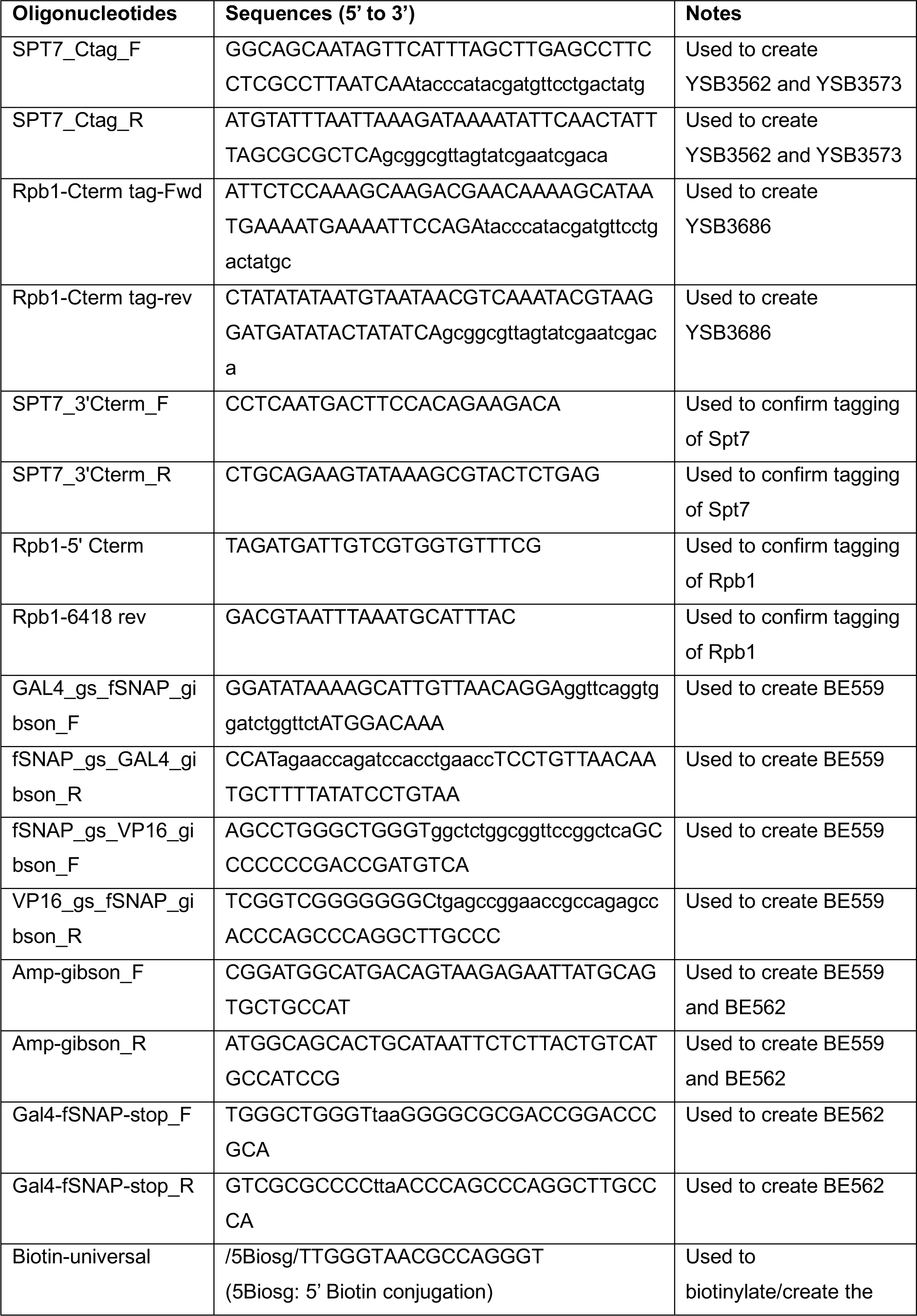
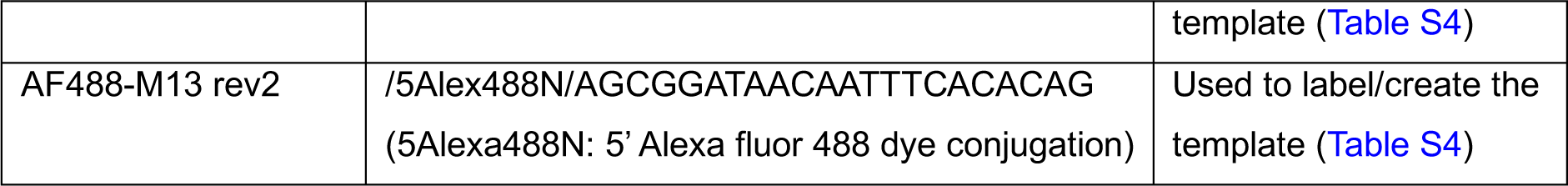
Oligonucleotides used in this study.

**Table S4:**
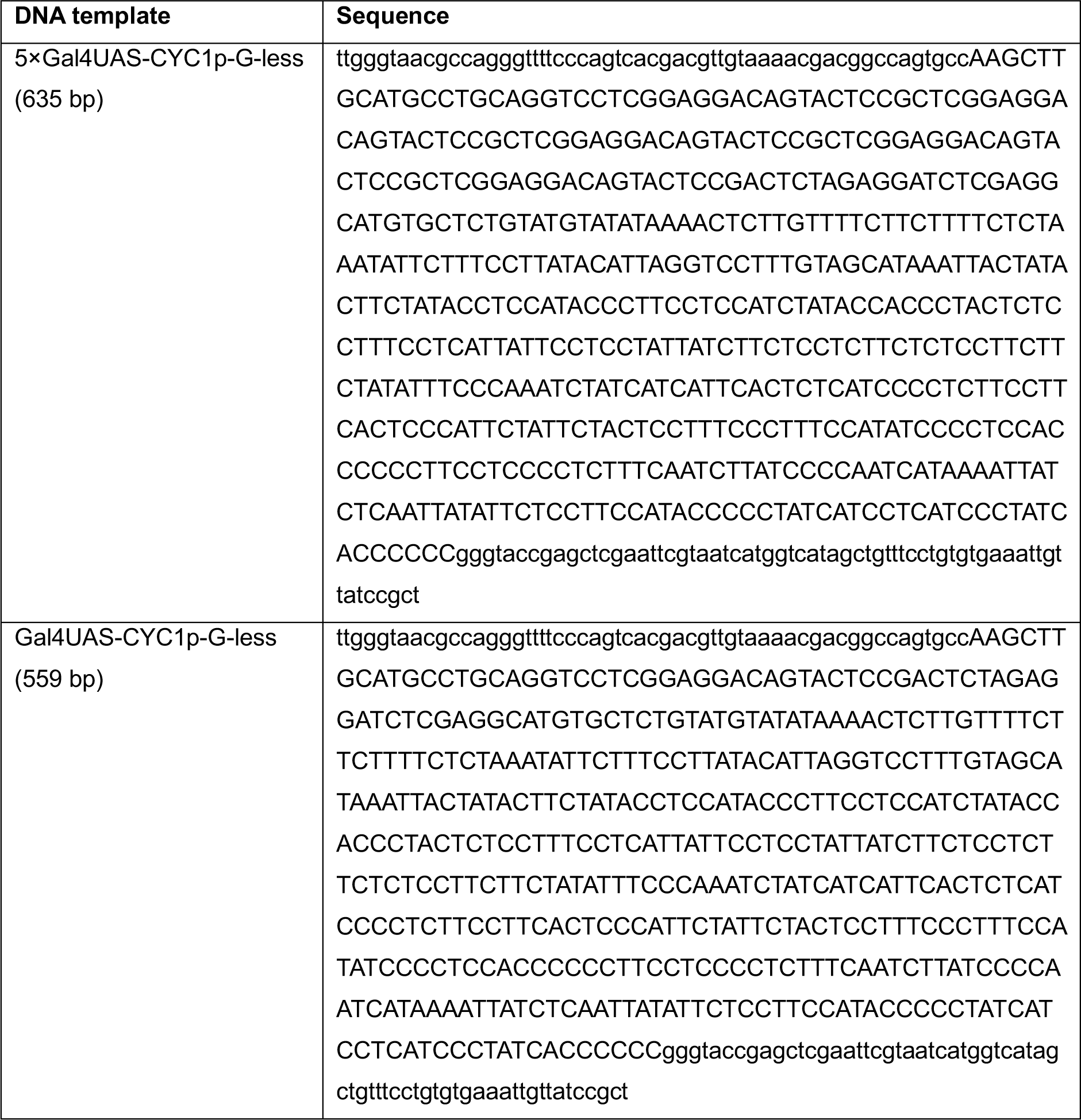
DNA templates.

**Table S5:** Kinetic parameters derived from fitting time-to-first binding curves for association of labeled factors with templates.

| Labeled factor<br>(Extract) | Fits<br>(Activator condition) | Template | $A_f$ | $k_{on} (s^{-1})$ | $k_{ns} (s^{-1})$ |
| --- | --- | --- | --- | --- | --- |
| Spt7-SNAP<sup>DY549</sup><br>(YSB3686) | Fig. 2B top and Fig. S3C<br>(10 nM Gal4-SNAP<sup>DY649</sup>-VP16) | Naked | $0.781 \pm 0.034$ | $2.79 \times 10^{-3} \pm 3.31 \times 10^{-4}$<br>(N=202) | $6.16 \times 10^{-5} \pm 1.53 \times 10^{-5}$<br>(N=195) |
| Spt7-SNAP<sup>DY549</sup><br>(YSB3686) | Fig. 2B bottom and Fig. S3C<br>(0 nM Activator) | Naked | $0.712 \pm 0.072$ | $1.46 \times 10^{-3} \pm 2.93 \times 10^{-4}$<br>(N=192) | $8.23 \times 10^{-5} \pm 1.83 \times 10^{-5}$<br>(N=197) |
| Spt7-SNAP<sup>DY549</sup><br>(YSB3686) | Fig. S3C<br>(3 nM Gal4-SNAP<sup>DY649</sup>-VP16) | Naked | $0.754 \pm 0.032$ | $2.68 \times 10^{-3} \pm 3.01 \times 10^{-4}$<br>(N=255) | $6.25 \times 10^{-5} \pm 1.55 \times 10^{-5}$<br>(N=203) |
| Spt7-SNAP<sup>DY549</sup><br>(YSB3686) | Fig. 4C<br>(30 nM Gal4-SNAP<sup>DY649</sup>-VP16) | Naked | $0.822 \pm 0.027$ | $5.25 \times 10^{-3} \pm 5.83 \times 10^{-4}$<br>(N=244) | $7.58 \times 10^{-5} \pm 1.74 \times 10^{-5}$<br>(N=179) |
| Spt7-SNAP<sup>DY549</sup><br>(YSB3686) | Fig. S3C<br>(0 nM Activator) | Chromatinized | $0.505 \pm 0.231$ | $6.62 \times 10^{-4} \pm 3.65 \times 10^{-4}$<br>(N=160) | $8.23 \times 10^{-5} \pm 1.83 \times 10^{-5}$<br>(N=197) |
| Spt7-SNAP<sup>DY549</sup><br>(YSB3686) | Fig. S3C<br>(3 nM Gal4-SNAP<sup>DY649</sup>-VP16) | Chromatinized | $0.598 \pm 0.150$ | $8.86 \times 10^{-4} \pm 3.02 \times 10^{-4}$<br>(N=167) | $6.25 \times 10^{-5} \pm 1.55 \times 10^{-5}$<br>(N=203) |
| Spt7-SNAP<sup>DY549</sup><br>(YSB3686) | Fig. S3C<br>(10 nM Gal4-SNAP<sup>DY649</sup>-VP16) | Chromatinized | $0.656 \pm 0.140$ | $8.49 \times 10^{-4} \pm 2.68 \times 10^{-4}$<br>(N=203) | $6.16 \times 10^{-5} \pm 1.53 \times 10^{-5}$<br>(N=195) |
| Spt7-SNAP<sup>DY549</sup><br>(YSB3686) | Fig. 4C<br>(30 nM Gal4-SNAP<sup>DY649</sup>-VP16) | Chromatinized | $0.643 \pm 0.049$ | $2.28 \times 10^{-3} \pm 3.93 \times 10^{-4}$<br>(N=155) | $7.58 \times 10^{-5} \pm 1.74 \times 10^{-5}$<br>(N=179) |
| Gal4- | Fig. S3C | Naked | NA | NA | NA |
| SNAP<sup>DY649</sup>_<br>VP16 | (0 nM<br>Activator) |  |  |  |  |
| Gal4-SNAP<sup>DY649</sup>_<br>VP16 | Fig. S3C<br>(3 nM Gal4-SNAP<sup>DY649</sup>_<br>VP16) | Naked | 0.451 ± 0.040 | 1.80 × 10<sup>-3</sup> ±<br>3.37 × 10<sup>-4</sup><br>(N=255) | 1.06 × 10<sup>-5</sup> ±<br>6.12 × 10<sup>-6</sup><br>(N=203) |
| Gal4-SNAP<sup>DY649</sup>_<br>VP16 | Fig. S3C<br>(10 nM Gal4-SNAP<sup>DY649</sup>_<br>VP16) | Naked | 0.774 ± 0.032 | 3.36 × 10<sup>-3</sup> ±<br>4.17 × 10<sup>-4</sup><br>(N=202) | 4.18 × 10<sup>-5</sup> ±<br>1.30 × 10<sup>-5</sup><br>(N=195) |
| Gal4-SNAP<sup>DY649</sup>_<br>VP16 | Fig. 4C<br>(30 nM Gal4-SNAP<sup>DY649</sup>_<br>VP16) | Naked | 0.842 ± 0.025 | 1.01 × 10<sup>-2</sup> ±<br>1.35 × 10<sup>-3</sup><br>(N=244) | 7.57 × 10<sup>-5</sup> ±<br>1.76 × 10<sup>-5</sup><br>(N=179) |
| Gal4-SNAP<sup>DY649</sup>_<br>VP16 | Fig. S3C<br>(0 nM<br>Activator) | Chromatinized | NA | NA | NA |
| Gal4-SNAP<sup>DY649</sup>_<br>VP16 | Fig. S3C<br>(3 nM Gal4-SNAP<sup>DY649</sup>_<br>VP16) | Chromatinized | 0.321 ± 0.187 | 9.29 × 10<sup>-4</sup> ±<br>4.60 × 10<sup>-4</sup><br>(N=167) | 1.06 × 10<sup>-5</sup> ±<br>6.12 × 10<sup>-6</sup><br>(N=203) |
| Gal4-SNAP<sup>DY649</sup>_<br>VP16 | Fig. S3C<br>(10 nM Gal4-SNAP<sup>DY649</sup>_<br>VP16) | Chromatinized | 0.557 ± 0.058 | 1.44 × 10<sup>-3</sup> ±<br>2.78 × 10<sup>-4</sup><br>(N=203) | 4.18 × 10<sup>-5</sup> ±<br>1.30 × 10<sup>-5</sup><br>(N=195) |
| Gal4-SNAP<sup>DY649</sup>_<br>VP16 | Fig. 4C<br>(30 nM Gal4-SNAP<sup>DY649</sup>_<br>VP16) | Chromatinized | 0.627 ± 0.044 | 5.08 × 10<sup>-3</sup> ±<br>1.04 × 10<sup>-3</sup><br>(N=155) | 7.57 × 10<sup>-5</sup> ±<br>1.76 × 10<sup>-5</sup><br>(N=179) |
| Gal4-SNAP<sup>DY649</sup>_<br>VP16 | Fig. S5B<br>(10 nM Gal4-SNAP<sup>DY649</sup>_<br>VP16) | Naked<br>(1xGal4<br>binding site<br>UAS) | 0.435 ± 0.073 | 1.52 × 10<sup>-3</sup> ±<br>4.19 × 10<sup>-4</sup><br>(N=140) | 7.75 × 10<sup>-5</sup> ±<br>1.21 × 10<sup>-5</sup><br>(N=368) |
| Spt7-SNAP<sup>DY549</sup><br>(YSB3562) | -<br>(30 nM Gal4-SNAP<sup>DY649</sup>_<br>VP16) | Chromatinized | 0.679 ± 0.159 | 1.11 × 10<sup>-3</sup> ±<br>4.49 × 10<sup>-4</sup><br>(N=127) | 5.83 × 10<sup>-5</sup> ±<br>1.28 × 10<sup>-5</sup><br>(N=329) |
| Spt7-SNAP<sup>DY549</sup> | -<br>(30 nM Gal4- | Chromatinized | 0.090 ± 0.272 | 3.23 × 10<sup>-3</sup> ±<br>4.50 × 10<sup>-2</sup> | 9.75 × 10<sup>-5</sup> ±<br>2.05 × 10<sup>-5</sup> |

| (YSB3562) | SNAP<sup>DY649</sup>) |  |  | (N=189) | (N=201) |
| --- | --- | --- | --- | --- | --- |
| Gal4-SNAP<sup>DY649</sup>-VP16 | -<br>(30 nM Gal4-SNAP<sup>DY649</sup>-VP16) | Chromatinized | 0.660 ± 0.054 | $5.58 \times 10^{-3} \pm 2.31 \times 10^{-3}$<br>(N=127) | $1.23 \times 10^{-4} \pm 1.88 \times 10^{-5}$<br>(N=329) |
| Gal4-SNAP<sup>DY649</sup> | -<br>(30 nM Gal4-SNAP<sup>DY649</sup>) | Chromatinized | 0.793 ± 0.038 | $3.67 \times 10^{-3} \pm 6.82 \times 10^{-4}$<br>(N=189) | $1.09 \times 10^{-4} \pm 2.28 \times 10^{-5}$<br>(N=201) |
| Spt7-HALO<sup>JF646</sup> (YSB3685) | Fig. S6A left<br>(200 nM Gal4-VP16) | Naked | 0.908 ± 0.024 | $9.69 \times 10^{-3} \pm 1.50 \times 10^{-3}$<br>(N=181) | $7.59 \times 10^{-5} \pm 1.28 \times 10^{-5}$<br>(N=356) |
| Spt7-HALO<sup>JF646</sup> (YSB3685) | Fig. S6A right<br>(200 nM Gal4-RAP1) | Naked | 0.895 ± 0.024 | $1.59 \times 10^{-2} \pm 2.22 \times 10^{-3}$<br>(N=172) | $1.07 \times 10^{-4} \pm 1.48 \times 10^{-5}$<br>(N=359) |
| Spt7-SNAP<sup>DY549</sup> (YSB3686) | Fig. 7A<br>(20 nM Gal4-SNAP<sup>DY649</sup>-VP16) | Chromatinized<br>(human unmodified) | 0.405 ± 0.041 | $2.31 \times 10^{-3} \pm 4.69 \times 10^{-4}$<br>(N=226) | $6.07 \times 10^{-5} \pm 9.68 \times 10^{-6}$<br>(N=478) |
| Spt7-SNAP<sup>DY549</sup> (YSB3686) | Fig. 7A<br>(20 nM Gal4-SNAP<sup>DY649</sup>-VP16) | Chromatinized<br>(human H4 tetra-actylated) | 0.572 ± 0.041 | $3.14 \times 10^{-3} \pm 4.50 \times 10^{-4}$<br>(N=189) | $6.07 \times 10^{-5} \pm 9.68 \times 10^{-6}$<br>(N=478) |
| Gal4-SNAP<sup>DY649</sup>-VP16 | Fig. 7A<br>(20 nM Gal4-SNAP<sup>DY649</sup>-VP16) | Chromatinized<br>(human unmodified) | 0.629 ± 0.041 | $2.47 \times 10^{-3} \pm 3.73 \times 10^{-4}$<br>(N=226) | $8.79 \times 10^{-5} \pm 1.20 \times 10^{-5}$<br>(N=478) |
| Gal4-SNAP<sup>DY649</sup>-VP16 | Fig. 7A<br>(20 nM Gal4-SNAP<sup>DY649</sup>-VP16) | Chromatinized<br>(human H4 tetra-actylated) | 0.738 ± 0.046 | $1.95 \times 10^{-3} \pm 3.80 \times 10^{-4}$<br>(N=189) | $8.79 \times 10^{-5} \pm 1.20 \times 10^{-5}$<br>(N=478) |
| Spt7-SNAP<sup>DY549</sup> (YSB3686) | Fig. S7A<br>(20 nM Gal4-SNAP<sup>DY649</sup>-VP16) | Chromatinized<br>(human unmodified) | 0.279 ± 0.043 | $3.30 \times 10^{-3} \pm 9.39 \times 10^{-4}$<br>(N=211) | $9.04 \times 10^{-5} \pm 1.59 \times 10^{-5}$<br>(N=279) |
| Spt7-SNAP<sup>DY549</sup> (YSB3686) | Fig. S7A<br>(20 nM Gal4-SNAP<sup>DY649</sup>-VP16) | Chromatinized<br>(human H4 tetra-actylated) | 0.348 ± 0.041 | $2.78 \times 10^{-3} \pm 6.13 \times 10^{-4}$<br>(N=273) | $9.04 \times 10^{-5} \pm 1.59 \times 10^{-5}$<br>(N=279) |
| Gal4-SNAP<sup>DY649</sup>-VP16 | Fig. S7A<br>(20 nM Gal4-SNAP<sup>DY649</sup>-VP16) | Chromatinized<br>(human unmodified) | 0.440 ± 0.044 | 2.47 × 10<sup>-3</sup> ± 5.83 × 10<sup>-4</sup><br>(N=211) | 6.11 × 10<sup>-5</sup> ± 1.27 × 10<sup>-5</sup><br>(N=279) |
| Gal4-SNAP<sup>DY649</sup>-VP16 | Fig. S7A<br>(20 nM Gal4-SNAP<sup>DY649</sup>-VP16) | Chromatinized<br>(human H4 tetra-actylated) | 0.559 ± 0.053 | 1.75 × 10<sup>-3</sup> ± 4.40 × 10<sup>-4</sup><br>(N=273) | 6.11 × 10<sup>-5</sup> ± 1.27 × 10<sup>-5</sup><br>(N=279) |
| Spt7-SNAP<sup>DY549</sup> (YSB3686) | Fig. 7E, no NTPs<br>(18 nM Gal4-SNAP<sup>DY649</sup>-VP16) | Naked | 0.889 ± 0.020 | 6.63 × 10<sup>-3</sup> ± 4.64 × 10<sup>-4</sup><br>(N=290) | 1.17 × 10<sup>-4</sup> ± 1.60 × 10<sup>-5</sup><br>(N=362) |
| Spt7-SNAP<sup>DY549</sup> (YSB3686) | Fig. 7E, + NTPs<br>(18 nM Gal4-SNAP<sup>DY649</sup>-VP16) | Naked | 0.895 ± 0.019 | 1.11 × 10<sup>-2</sup> ± 8.86 × 10<sup>-4</sup><br>(N=321) | 1.69 × 10<sup>-4</sup> ± 2.23 × 10<sup>-5</sup><br>(N=281) |
| Spt7-SNAP<sup>DY549</sup> (YSB3686) | Fig. 7E, no NTPs<br>(18 nM Gal4-SNAP<sup>DY649</sup>-VP16) | Chromatinized | 0.763 ± 0.225 | 4.64 × 10<sup>-4</sup> ± 3.28 × 10<sup>-4</sup><br>(N=178) | 1.17 × 10<sup>-4</sup> ± 1.60 × 10<sup>-5</sup><br>(N=362) |
| Spt7-SNAP<sup>DY549</sup> (YSB3686) | Fig. 7E, + NTPs<br>(18 nM Gal4-SNAP<sup>DY649</sup>-VP16) | Chromatinized | 0.541 ± 0.070 | 1.91 × 10<sup>-3</sup> ± 4.88 × 10<sup>-4</sup><br>(N=165) | 1.69 × 10<sup>-4</sup> ± 2.23 × 10<sup>-5</sup><br>(N=281) |
| Gal4-SNAP<sup>DY649</sup>-VP16 | Fig. 7E, no NTPs<br>(18 nM Gal4-SNAP<sup>DY649</sup>-VP16) | Naked | 0.867 ± 0.022 | 1.02 × 10<sup>-2</sup> ± 1.19 × 10<sup>-3</sup><br>(N=290) | 8.95 × 10<sup>-5</sup> ± 1.53 × 10<sup>-5</sup><br>(N=362) |
| Gal4-SNAP<sup>DY649</sup>-VP16 | Fig. 7E, + NTPs<br>(18 nM Gal4-SNAP<sup>DY649</sup>-VP16) | Naked | 0.943 ± 0.003 | 1.45 × 10<sup>-2</sup> ± 1.36 × 10<sup>-3</sup><br>(N=321) | 8.93 × 10<sup>-5</sup> ± 1.53 × 10<sup>-5</sup><br>(N=281) |
| Gal4- | Fig. 7E, no | Chromatinized | 0.490 ± 0.050 | 2.37 × 10<sup>-3</sup> ± | 8.95 × 10<sup>-5</sup> ± |
| SNAP<sup>DY649</sup>-<br>VP16 | **NTPs**<br>(18 nM Gal4-SNAP<sup>DY649</sup>-<br>VP16) | | | $5.24 \times 10^{-4}$<br>(N=178) | $1.53 \times 10^{-5}$<br>(N=362) |
| Gal4-SNAP<sup>DY649</sup>-<br>VP16 | **Fig. 7E, +**<br>**NTPs**<br>(18 nM Gal4-SNAP<sup>DY649</sup>-<br>VP16) | Chromatinized | $0.635 \pm 0.045$ | $2.81 \times 10^{-3} \pm$<br>$4.62 \times 10^{-4}$<br>(N=165) | $8.93 \times 10^{-5} \pm$<br>$1.53 \times 10^{-5}$<br>(N=281) |
| Spt7-SNAP<sup>DY549</sup><br>(YSB3686) | **Fig. S7C**<br>**middle**<br>(25 nM Gal4-SNAP<sup>DY649</sup>-<br>VP16) | Chromatinized<br>(human<br>unmodified) | $0.455 \pm 0.037$ | $3.89 \times 10^{-3} \pm$<br>$5.67 \times 10^{-4}$<br>(N=211) | $5.15 \times 10^{-5} \pm$<br>$9.91 \times 10^{-6}$<br>(N=402) |
| Spt7-SNAP<sup>DY549</sup><br>(YSB3686) | **Fig. S7C**<br>**middle**<br>(25 nM Gal4-SNAP<sup>DY649</sup>-<br>VP16) | Chromatinized<br>(human H3<br>tetra-acetylated) | $0.443 \pm 0.035$ | $5.76 \times 10^{-3} \pm$<br>$1.19 \times 10^{-3}$<br>(N=267) | $5.15 \times 10^{-5} \pm$<br>$9.91 \times 10^{-6}$<br>(N=402) |
| Gal4-SNAP<sup>DY649</sup>-<br>VP16 | **Fig. S7C left**<br>(25 nM Gal4-SNAP<sup>DY649</sup>-<br>VP16) | Chromatinized<br>(human<br>unmodified) | $0.525 \pm 0.044$ | $4.57 \times 10^{-3} \pm$<br>$1.83 \times 10^{-3}$<br>(N=211) | $8.43 \times 10^{-5} \pm$<br>$1.27 \times 10^{-5}$<br>(N=402) |
| Gal4-SNAP<sup>DY649</sup>-<br>VP16 | **Fig. S7C left**<br>(25 nM Gal4-SNAP<sup>DY649</sup>-<br>VP16) | Chromatinized<br>(human H3<br>tetra-acetylated) | $0.569 \pm 0.033$ | $5.80 \times 10^{-3} \pm$<br>$1.15 \times 10^{-3}$<br>(N=267) | $8.43 \times 10^{-5} \pm$<br>$1.27 \times 10^{-5}$<br>(N=402) |

**\*A<sub>f</sub> is the active fraction of template molecules.**
**k<sub>on</sub> and k<sub>ns</sub> are the apparent first-order rate constants of binding factors on- and off-target (non-specific) sites.**
**N is the total number of on- or off-template sites.**
**Standard errors of fit parameters were determined by bootstrapping with 2000 iterations.**

